# Inhibiting ezrin triggers formin-mediated actin remodelling reducing cellular mechano-protection

**DOI:** 10.1101/2025.07.22.666169

**Authors:** Ananya Biswas, Rinku Kumar, T N Husain, Jibitesh Das, Amrutamaya Behera, Bidisha Sinha

## Abstract

Ezrin plays a crucial role in linking the cortical actin cytoskeleton to the plasma membrane in animal cells. Inactivated by small molecule ezrin inhibitors (EzrInh), ezrin is known to regulate cellular functions like adhesion and motility. However, whether ezrin plays any significant role in protecting cells against mechanical stresses is less explored. We use hypo-osmotic shock (HS) to challenge cells and study the mechano-protective role of ezrin. Inhibition of ezrin phosphorylation led to increased cell rupture rates on hypo-shock, with a greater impact at later time points (∼ 10 min). The time scales match with that of enhanced tension contrast at edges at the basal plane. We also observed a lowered relative retraction rate at protrusions created by hypo-shock and a less contractile mid-plane cortex – both indicate the role of a weakening of the cortex by EzrInh in compromising membrane integrity on hypo-shock. Concomitantly, EzrInh also increased stress fibres (SFs), traction forces and cell spreading clearly indicative of additional cellular pre-stress. Utilizing formin inhibitor (SMIFH2), we showed that formin-mediated actin polymerization was critical for the SF response to EzrInh while it also prevented the subsequent EzrInh-treatment from enhancing cell rupture rates. Together EzrInh reduced mechano-protection in cells by tilting of balance of actin’s organization from the cortex to SF enhancing formin-mediated SF assembly.

**Highlights:**

- Hypo-osmotically shocked CHO cells rupture more on ezrin inhibition
- Ezrin inhibition weakens contractility of cortical actin but enhances stress fibre numbers at the base
- Hypo-osmotic shock creates basal protrusions that are also retracted.
- Retractive potential in cells is weakened by ezrin inhibition
- Formin inhibitor prevents net increase in stress fibres by inhibition of ezrin
- On pre-treatment with formin inhibitor, subsequent inhibition of ezrin doesnot render cells more vulnerable to hypo-osmotic shock

## Introduction

Ezrin is a key member of the ERM (ezrin-radixin-moesin) family of proteins that serve as linkers between the plasma membrane and the actin cytoskeleton. Ezrin facilitates actin attachment to the membrane and, upon phosphorylation, stabilizes this interaction, thereby contributing to cell shape and mechanical resilience. Ezrin is present in an inactive state in the cytoplasm and requires either binding of its head domain (N-ERMAD) to PIP2 at cell membrane or phosphorylation at Thr567 position for activation ^1^. This unique ability to link membrane proteins to the actin cytoskeleton makes ezrin a central player in mediating signal transduction across a number of pathways such as Rho GTPase signalling, EGF-EGFR, PI3K/AKT pathway, etc ^2–5^ .

Cells constantly face mechanical stress^2^ —from shear forces in blood flow to tension in alveoli and loads on joints which are sensed and transduced^6^. During metastasis cancer cells migrate through varied mechanical landscape and need mechanisms to protect themselves^7^. Interestingly, although ezrin’s overexpression is not a hallmark of metastasis, it is known to facilitate metastatic dissemination in multiple cancers. While its role has been addressed primarily from a signalling perspective – very little is known about its contribution to cell mechanics which too could potentially impact metastasis. Ezrin inhibition affects membrane tension^8^ but whether it determines cell survival under stress is less studied. The actomyosin network underlying the membrane serves as the first line of defence by ascribing enhanced stiffness and hence resilience to the cell surface^9,10^. However, to offer this protection it must remain attached to membrane – via ezrin. Being a contractile material, in the absence of actin-membrane linkage the network is expected to shrink away from the membrane. To understand if ezrin serves as a mechano-protector, cell survival in the face of mechanical stress needs to be studied under control and ezrin-inhibited conditions.

Hypo-osmotic shock mechanically perturbs cells by causing water influx, leading to cell swelling and increased membrane stress. This stretch activates mechanosensitive pathways, disrupts cytoskeletal organization, and can induce regulatory mechanisms (like caveolae flattening^11^, membrane rupture or repair responses. It, therefore, serves as a valuable tool to study cellular mechano-resilience and membrane-cytoskeleton interactions under stress. Hypo-osmotic stress in some cells are known to induce stress fibre formation. Stress fibres are the major tension-generating and load-bearing mechanosensitive structures, composed of bundles of F-actin along with myosin IIa and other accessory proteins^12^. Stress fibres can be classified into a number of sub-types: perinuclear cap fibre, dorsal stress fibre, transverse arc, and ventral stress fibre, cortical stress fibres etc. based on their localization and associated proteins ^13–15^. The peripheral stress fibres are devoid of myosin IIa and located at the periphery of the cell. They are involved in generating the protruding and non-protruding edges of a migrating cell ^16^. The cortical stress fibres, on the other hand, are present just beneath the plasma membrane. Interspersed with myosin IIa, these are involved in generating contractile forces ^17^. The stress fibre generation involves Rho signalling and could either be stabilized by the ROCK pathway involving myosin or by stabilizing nucleation by formins. Studies have shown formins to be inherently mechanosensitive ^18,19^ - but whether involved in ezrin’s mechano-protective role is not known.

In this work, we use hypo-osmotic shock to challenge cells and score for resilience. Ezrin inhibitor is used to evaluate the role of ezrin in providing resilience against hypo-shock. Using Interference Reflection Microscopy (IRM) and fluorescence lifetime imaging (FLIM) we evaluate the impact on the membrane mechanics by inhibiting ezrin or administering hypo-shock. Mapping traction forces reveals how cells not only show a direct effect of inhibition of ezrin but respond strongly by changing their mechanical state. Stress fibre enhancement by EzInh and dynamics of the cell edge on hypo-shock are captured by following cells using Total Internal Reflection Fluorescence (TIRF) Microscopy expressing a F-actin binding fluorescently-tagged peptide Lifeact-mCherry. With combination of formin, ROCK and ezrin inhbitors, this work unravels how ezrin and formin together maintain the intricate balance of cortical actin and stress fibres from the same pool of F-actin to transit the cell to a less dynamic and more vulnerable static state on ezrin inhibition.

## Results

### Ezrin Inhibition enhances rupture on Hypo-Osmotic Shock

Previous studies have shown hypo-osmotic shock to challenge membrane integrity^11,20^. Hence, in order to understand cellular mechano-protection, we designed a hypo-osmotic shock-based rupture assay which would enable us to trigger mechano-protective responses in cells on being subjected to a global stressor^21^. For this, we first chose to study the effect of inactivating ezrin in cells followed by hypo-osmotically shocking them to elucidate the role of ezrin in conferring mechanical protection. Ezrin Inhibitor (EzrInh) was used at 40 μM concentration for 1 hr followed by assaying cells for ezrin’s mechano-protective role. We first established EzrInh to reduce the levels of P-ezrin at the basal plane by Total Internal Fluorescence Microscopy (TIRF) (**Fig. 1A, Fig. S1A**) and in whole cells by Flow cytometry (**Fig. S1B**).

**Figure 1.**
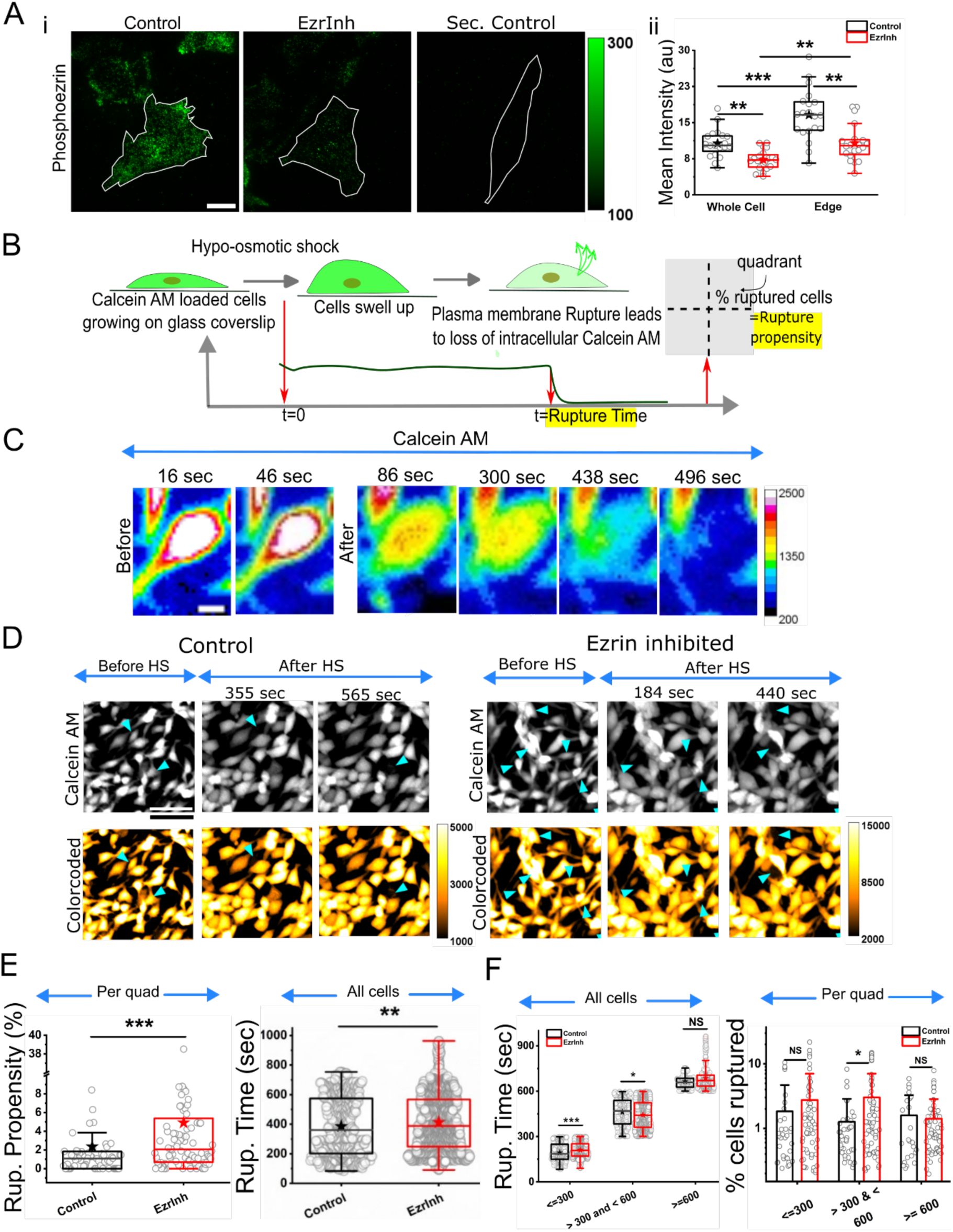
Rupture assay of ezrin inhibited CHO cells upon hypo-osmotic shock treatment. A. Quantification of phosphor-ezrin (p-ezrin) upon adding EzrInh (40 µM, 1 hr.) to CHO cells. i. Representative TIRF images of immunofluorescent p-ezrin at basal plane of cells. Scale: 10 µm. ii. Comparison of p-ezrin intensity upon EzrInh at basal plane versus only at cell periphery (edge). N= 2 independent experiments, n_control_ = 40 cells, n_EzrInh_ = 28 cells. B. Graphical representation of hypo-osmotic shock mediated rupture assay in CHO cells. C. Heatmap showing change in Calcein AM intensity in a cell undergoing rupture. Scale: 5 µm. D. Representative epi-fluorescent images of Calcein AM (2.5 µM, 30 min) loaded CHO cells before and after hypo-osmotic shock upon EzrInh treatment. Rupture events denoted by cyan arrows. Scale: 50 µm. E. Comparison of rupture propensity and rupture time upon EzrInh treatment. Rupture propensity is plotted quadrant-wise whereas rupture time of all rupturing cells are plotted. F. Comparison of rupture time across three time scales. Also shown is a quadrant-wise comparison of percent of cells rupturing over the same time scales. N= 9 experimental repeats. No. of quadrants: Q_Control_ = 64, Q_EzrInh_ = 92; Total no. of ruptures: nrup_Control_ = 487, nrup_EzrInh_ = 1364. Statistical significance was determined by Mann Whitney U-test where *p < 0.05, **p < 0.01, ***p < 0.001 and ns p > 0.05.

Next, in order to assay mechano-protection, ezrin inhibited cells were loaded with Calcein AM, a tracer dye, at concentrations of 2.5 µM followed by hypo-osmotically (95% shock) shocking the cells. On following the cells for 10 min and imaging at every 2 sec interval, events of single cell ruptures could be identified (**Fig. 1B, C**). We have defined two parameters, namely, Rupture Propensity (percentage cells thar rupture that rupture in a given area – usually a quadrant of the imaging area corresponding ∼ 330 μm x 330 μm and Rupture time (time taken by a cell to rupture from the time of administration of hypo-shock) to understand loss of mechano-protection (**Fig. 1B, Fig. S2**). In general cells are highly resilient to hypo-osmotic shock with only ∼ 2% cells rupturing within the first 10 min. On ezrin inhibition, cells showed an increase in the rupture propensity indicating a loss of mechano-protection. However, in the whole pool, the median time taken by ezrin inhibited cells to rupture showed an increased (**Fig 1D, E**). From the histogram of rupture times (**Fig. S3 A-B**), a bimodal distribution emerged indicating cells’ response to ezrin inhibition may be dependent on a range of timescales.

Therefore, we further classified the rupture times into three different timescales: fast (≤ 300 sec), intermediary (>300 sec & <600 sec) and slow (≥ 600 secs). Also, upon comparing the percentage of cells rupturing in the above-mentioned timescales quadrant wise and comparing the time taken by these cells to rupture at the respective timescales, an interesting trend emerged. The mechano-protective role of ezrin could be felt more at the intermediary timescales (∼5-10 mins) corroborated by the increase in percentage of rupturing cells at this timescale (**Fig. 1F**).

This establishes that in CHO cells ezrin has a mechano-protective role. However, how and why the ruptures were more pronounced at an intermediate timescale still required clarity.

### Altered membrane mechanical response in cells pre-treated with EzrInh

In order to understand how ezrin inhibition lead to a loss of mechano-protection, we next looked at the impact of ezrin inhibition on the actomyosin cortex. For this, we first probed into the dynamics of myosin IIa upon ezrin inhibition as myosin IIa is involved in conferring contractility to the actomyosin cortex^22^. Using Fluorescence Recovery upon Photobleaching (FRAP), we determined the turnover rate of fluorescently-tagged myosin IIa upon ezrin inhibition. Slow turnover of myosin IIa is known to correlate with enhanced contractile tension^22,23^. We observed that on ezrin inhibition, cells exhibited a faster turnover rate indicating a weaker actomyosin cortex (**Fig. 2A**). However, it was still unclear whether weakening of actomyosin cortex could lead to increased membrane ruptures.

**Figure 2.**
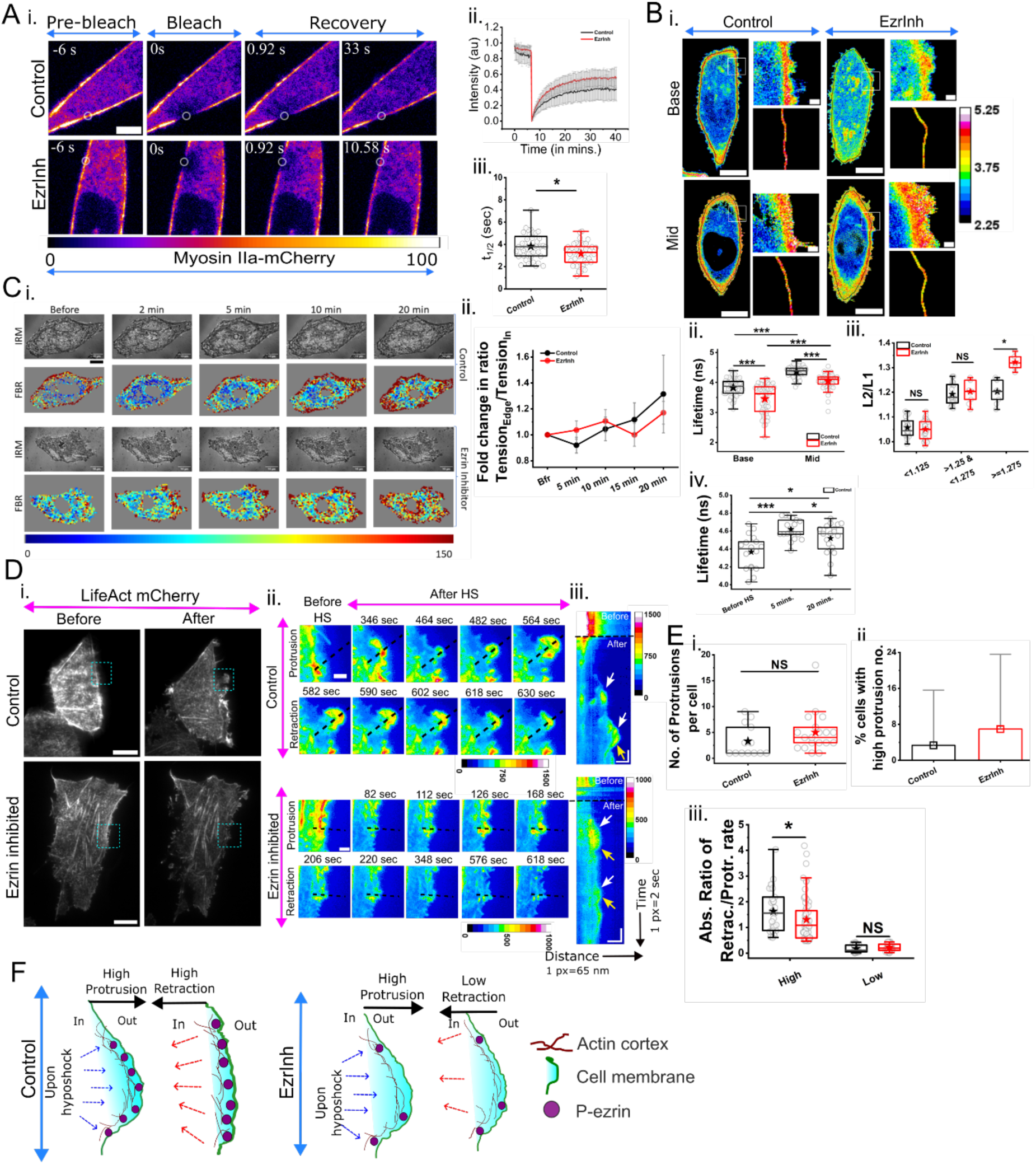
Actomyosin and membrane remodelling upon ezrin inhibition and hypo-osmotic shock. A. i. Representative images of fluorescence recovery in CHO cells transiently expressing myosin IIA-mCherry. ii. Comparison of recovery characteristics and half-life of myosin IIa-mCherry upon ezrin inhibition. Scale: 5 µm. N= 3 independent experiments, n_Control_ = 40 cells, n_EzrInh_ = 42 cells. B. i. Representative FLIM images captured at the basal as well as mid-plane of CHO cells stained with Flipper-TR (1 µM, 50 min). Scale: 10 µm. Zoomed-in images show a part of the cell membrane (*top*) along with the corresponding ROI of 2-pixel thickness (*bottom*) used for analysis. Scale: 1 µm. ii. Comparisons of lifetimes at base and mid plane upon ezrin inhibition. iii. Comparison of ratio of lifetimes (L2/L1) on EzrInh treatment. iv. Fluorescence lifetime increases upon hypo-osmotic shock. N= 3 independent experiments, n_Control_ = 28 cells, n_EzrInh_ = 30 cells. v. Ratio of lifetimes at mid plane (L2) and base (L1). N=3 independent experiments, n_Control_ = 57 cells, n_EzrInh_ = 57 cells. iv. C. i. Representative IRM images of CHO cells with corresponding FBR maps of EzrInh CHO cells upon hypo-osmotic shock. Scale: 10 µm. ii. Comparison of fold change in ratio of tension at cell edge versus inside. N=4 independent experiments, n_Control_ = 23 cells, n_EzrInh_ = 24 cells with same cells followed over time. Error bars represent standard errors. D. Representative TIRF images of CHO cells transiently expressing LifeAct mCherry before and after hypo-osmotic shock. Scale: 10 µm. ii. Zoomed-in view showing actin dynamics when hypo-osmotic shock is administered on ezrin-inhibited cells. Scale: 2 µm. iii. Kymograph generated across line ROI (black dotted line in ii) represent evolution of protrusions (white arrows) and corresponding retractions (yellow arrows). X-axis represents distance across ROI (1 pixel= 65 nm) while time on Y-axis (1 pixel= 2 sec). Scale: 2 µm (X-axis), 60 sec (Y-axis). E. i. Comparison of number of protrusions per cell. ii. Quantification of percentage of cells showing high protrusion numbers (>8 protrusions per cell). iii. Comparison of absolute ratio of rate of retraction to rate of protrusion. N= 6 independent experiments, n_Control_ = 15 cells, n_EzrInh_ = 20 cells. Statistical significance was determined by Mann Whitney U-test where *p < 0.05, **p < 0.01, ***p < 0.001 and ns p > 0.05.

To understand if the cell membrane dynamics displayed signatures of weakening, we next assayed membrane mechanics using Fluorescence Lifetime Microscopy (FLIM) and Interference Reflection Microscopy (IRM). Since, cortex contractility was evaluated at the mid-planes of cells, FLIM was used to measure the change in tension upon hypo-osmotic shock. Towards this, cells were incubated with FliptR (Fluorescent LIPid Tension Reporter) at a concentration of 1 µM^24^ and their fluorescence lifetime measured before and after hypo-osmotic shock (**Fig. 2B i**). Previous studies indicate that although fluorescence lifetime of FliptR (a membrane-integrating molecule) is affected by lipid compaction, it can be used as a tension change reporter if crosschecked with other techniques. Ezrin inhibition resulted in a reduction in fluorescence lifetime at both the basal as well as mid-planes of cells (**Fig. 2B ii**). However, the ratio of mid-to-basal plane lifetime collated measured for single cells showed that in resting condition (without hypo-shock), ezrin inhibition altered the population such that the cells with high tension contrast (mid to base ratio >= 1.275) had a stronger contrast (**Fig. 2B iii**). Hypo-osmotic shock transiently enhanced lifetime ∼ 5 min after which it started reducing for control (**Fig. 2B iii**) and ezrin-inhibited cells (**Fig. S4C**). The mid to base ratio increased on hyposhock (**Fig. S4D**) although the change was similar for control and ezrin-inhibited cells. Since we did not record any excessive rise in tension for the whole cells but an enhanced contrast in ezrin-inhibited cells, we next studied the evolution of any tension contrast at the basal plane.

We used IRM to measure nanometric height fluctuations of the basal membrane (**Fig. 2C**). From the power spectral density (PSD), an effective tension of the basal cell surface was estimated. We followed single cells through time upon hypo-osmotic shock and mapped the effective tension at regions where the membrane is expected to be within ∼ 100 nm (termed FBRs-first branch regions). FBRs were also segregated based on their position (**Fig. S4A**). When averaged values over whole basal membrane were compared through hypo-osmotic shock, the data revealed no mechanically drastic effect on ezrin-inhibited cells (**Fig. 2C i, S4B**). Next we measured, for every cell, the ratio of the tension at the cell edge (from FBRs which were with 10 pixels of the cell boundary) and that of other regions (**Fig. S4A**). The fold change in this tension ratio was noted for every cell. The averaged fold change was compared between EzInh and Control conditions. We observed that the difference between EzrInh and Control cells was maximized at the intermediary timescale (**Fig. 2C ii**)– as observed in the rupture timescale distribution (**Fig. 1E**).

Together, we demonstrated that on hypo-shock, ezrin inhibited cells get more mechanically challenged by being more heterogeneous at the intermediate time points. This could indicate that only certain localized regions might be more affected. Hence, we next investigated regions where hypo-osmotic shock resulted in outward protrusions at the basal plane of the cell.

### EzrInh pretreatment causes hypo-osmotically stressed cells to display higher protrusion rates and lowered retraction rates at edges

In order to understand the same, we decided to probe into the dynamics of actin remodelling upon hypo-osmotic shock. Using Total Internal Fluorescence (TIRF) microscopy, actin dynamics upon hypo-osmotic shock in ezrin inhibited cells transiently expressing LifeAct-mCherry was recorded for 10 min with an interval of 2 sec. From the time series, cells were shown to produce protrusions (**Fig. 2 D i-iii, Fig. S5**) in response to hypo-osmotic shock. Although the data suggested that EzrInh enhanced the total number of protrusions per cell with a greater percentage of cells with >8 protrusions emerging after hypo-shock, the differences were not statistically significant (**Fig. 2E – top and middle**). We measured retraction and protrusion rates from kymographs (**Fig. 2Diii, Fig. S5A, B**) and found the ratio of retraction to protrusion rate. Above a threshold (∼ 0.45), the average retraction-to-protrusion rate ratio for ezrin-inhibited cells showed a significant decrease than control cells (**Fig. 2E bottom**). The threshold was used because the distribution of this ratio indicated two populations (**Fig. S5C**). This lowering of the retraction-to-protrusion rate may indicate that reduction in p-ezrin due to Ezrinh hampered the actomyosin network’s ability to exert enough pulling forces as opposed to the hypo-osmotic induced swelling pushing on the membrane. The imbalance in rates could lead to net outward force buildup in certain regions of the membrane promoting cell ruptures (**Fig. 2F**).

Continuing investigations about the basal actin we asked whether the loss of cortical contractility was part of the whole basal plane losing contractility or was it especially the membrane adjoining cortical actin where this effect was evident.

### EzrInh-treatment enhances stress fibres and traction force

While our previous studies focussed on the effect of hypo-osmotic shock, here we assessed the state of basal actin in cells on ezrin inhibition. CHO cells in either control or ezrin-inhibited conditions were fixed and F-actin labelled with fluorescently-tagged Phalloidin and the basal cell surface imaged by TIRF (**Fig. 3A top, Fig. S6**). Clear pattern of stress fibres (SFs) emerged from these images. Images were analyzed by removing slow-varying background from inside the cell (**Fig. 3A bottom**) such that SFs could be clearly distinguished even when weakly intense from their immediate background. Cells in control and ezrin-inhibited conditions were further compared (**Fig. 3B i**) and it was observed that the total number of SFs increased on ezrin inhibition while the cell spread area also increased (**Fig. 3B ii**). Since the distribution of stress fibres is expected to change with changing cell shape, to reducing cell shape variability in the population, we used substrate micropatterning^25^ (**Fig. 3C i**) to restrict cell spreading to line shaped patterns (cell-adhesive line of width 10 μm and 15 μm spacing between lines) in this section of the work.

**Figure 3.**
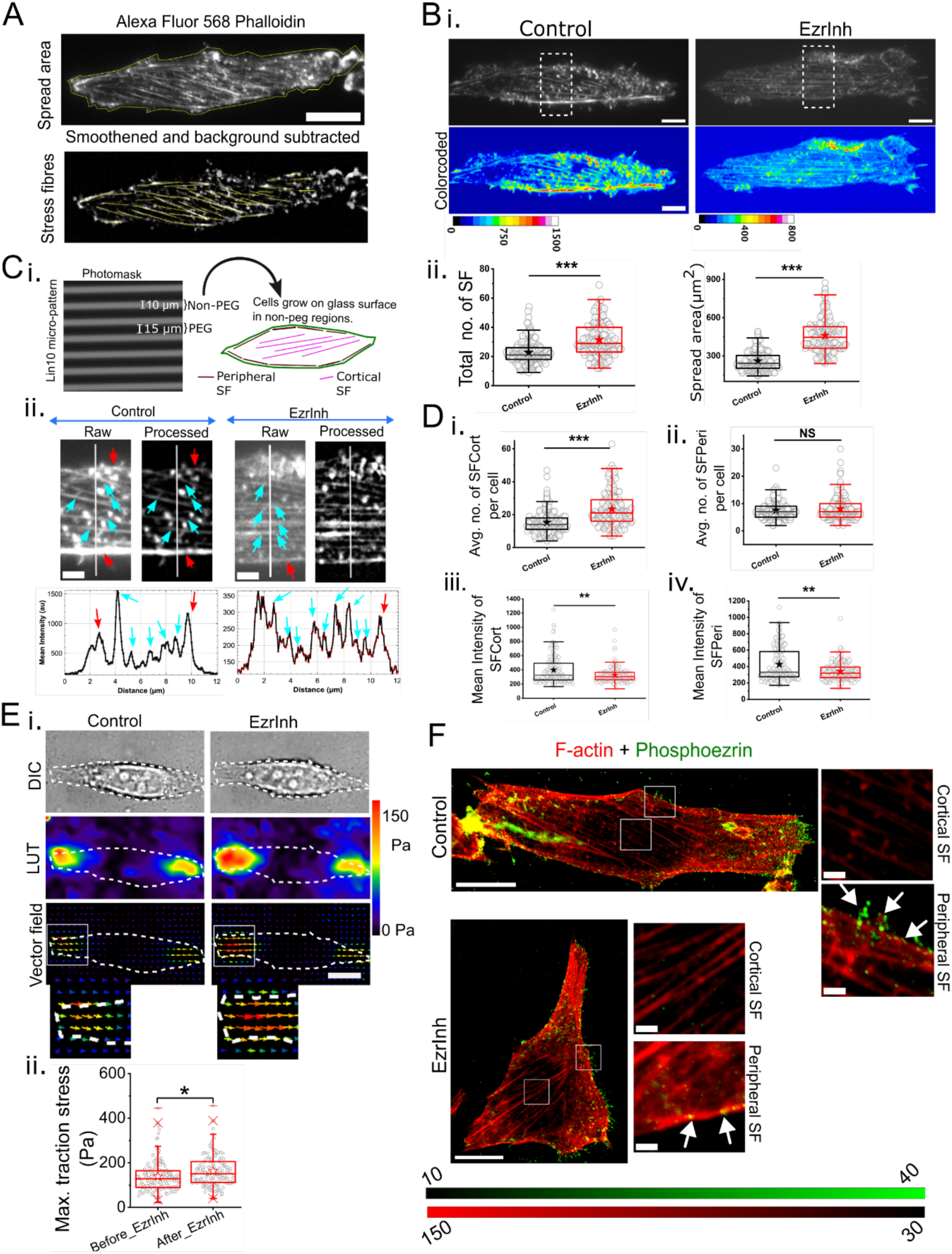
Ezrin inhibition enhances stress fibres and increases traction forces. A. Representative TIRF images of F-actin labelled with Alexa Fluor 568 PhalloidinLin10 micropatterned CHO cell. Stress fibres labelled with Alexa Fluor 568 Phalloidin. Cell spread area (*top*) and stress fibres (*bottom*) are defined by yellow ROIs. Scale bar: 10 µm. B. i. Representative TIRF images of stress fibres on ezrin inhibition along with corresponding heatmap where F-actin intensity is denoted by associated colorbar. ii. Comparison of total number of stress fibres and spread area of cells on EzrInh treatment. N= 3 independent experiments, n_Control_=169 cells, n_EzrInh_=175 cells. C. i. Graphical representation of micropatterning and distribution of peripheral stress fibres (*in maroon*) and cortical stress fibres (*in magenta*) at the basal plane. ii. Zoomed-in portion of cell showing cortical (*cyan arrows*) and peripheral (*red arrows*) stress fibres in ezrin-inhibited cells. (*left*) raw image and (*right*) smoothened and background subtracted for better visualization of stress fibres. Scale: 2 µm. Also shown is an intensity profile (*bottom*) of peripheral and cortical stress fibres across line ROI (*top*). D. i, ii. Comparison of average number of peripheral (SFPeri) and cortical stress fibres (SFCort). iii, iv. Comparison of mean intensity of SFPeri and SFCort. E. Traction force microscopy to measure effect of ezrin inhibition on traction stress of cells. i. Representative images of Lin10 micropatterned cells. Vector field image shows displacement of fluorescent beads from which traction stress is calculated. ii. Comparison of maximum traction stress on EzrInh treatment. N= 2 independent experiments. n_Control_ = 152 cells, n_EzrInh_ = 152 cells. F. Super-resolution images of F-actin and p-ezrin at the basal plane on ezrin inhibition. Zoomed-in view shows p-ezrin in stress fibre and peripheral region (*white arrows*). Statistical significance was determined by Mann Whitney U-test where *p < 0.05, **p < 0.01, ***p < 0.001 and ns p > 0.05, with Bonferroni correction for stress fibre analysis.

With similar shapes for all cells, we next separated SFs based on their location. SFs at the cell periphery were counted as peripheral SFs and others counted as cortical SFs – in line with previous studies (**Fig. 3C i**). We observed that on ezrin inhibition more number of finer SFs (corresponding to cortical SFs) could be observed in the cell (**Fig. 3C ii**). On quantification, it could be established that number of cortical SFs and no peripheral SFs were enhanced on ezrin-inhibition (**Fig. 3D i-ii**). The ratio of cortical to peripheral SF calculated for single cells also showed an increase (**Fig. S7A left**). Although the mean intensity of both cortical and peripheral SFs has decreased (**Fig. 3D iii-iv**) their ratio calculated in single cells remained unchanged (**Fig. S7A right**). The increase in number of SFs was supported by an increase in maximum and average traction forces as well (**Fig. 3E, Fig. S7B**) validating the functional significance of the enhanced SFs. The functional significance was further supported by ezrin-inhibition slowly wound closure in scratch assay (**Fig. S7B**). It is important to note that the p-ezrin itself overlaps very little with cortical stress fibres although it has some overlap with peripheral stress fibres supporting the possibility of their balance being disrupted by ezrin inhibition (**Fig. 3E**).

### Formin Dependency in Ezrin-inhibited Cells

Previous studies show stress fibre generation is mediated by either ROCK or formins. Hence, in order to understand whether the EzrInh-mediated increase in stress fibres was also through ROCK or formin pathway, we looked into stress numbers upon ezrin-inhibition in ROCK or formin inhibited cells. Using inhibitors for ROCK (Y-27632 or ROCKInh), and formin inhibitor, **SMIFH2,** for pre-treating cells, we calculated the total number of SFs upon subsequent ezrin inhibition (**Fig. 4A, Fig. S8**). Note that the comparison uses both pools with ROCKInh/SMIFH2 pre-treatment. The effect of the pre-treatments alone varied (**Fig. S9**). We observed that ezrin-inhibition in ROCKInh pre-treated cells increased cell spread area (**Fig. 4B i**) and total number of SFs (**Fig. 4B ii**). The number cortical SFs were enhanced (**Fig. 4B iii**) and not peripheral SFs (**Fig. 4B iii**) as was observed on inhibiting ezrin in control cells.

**Figure 4.**
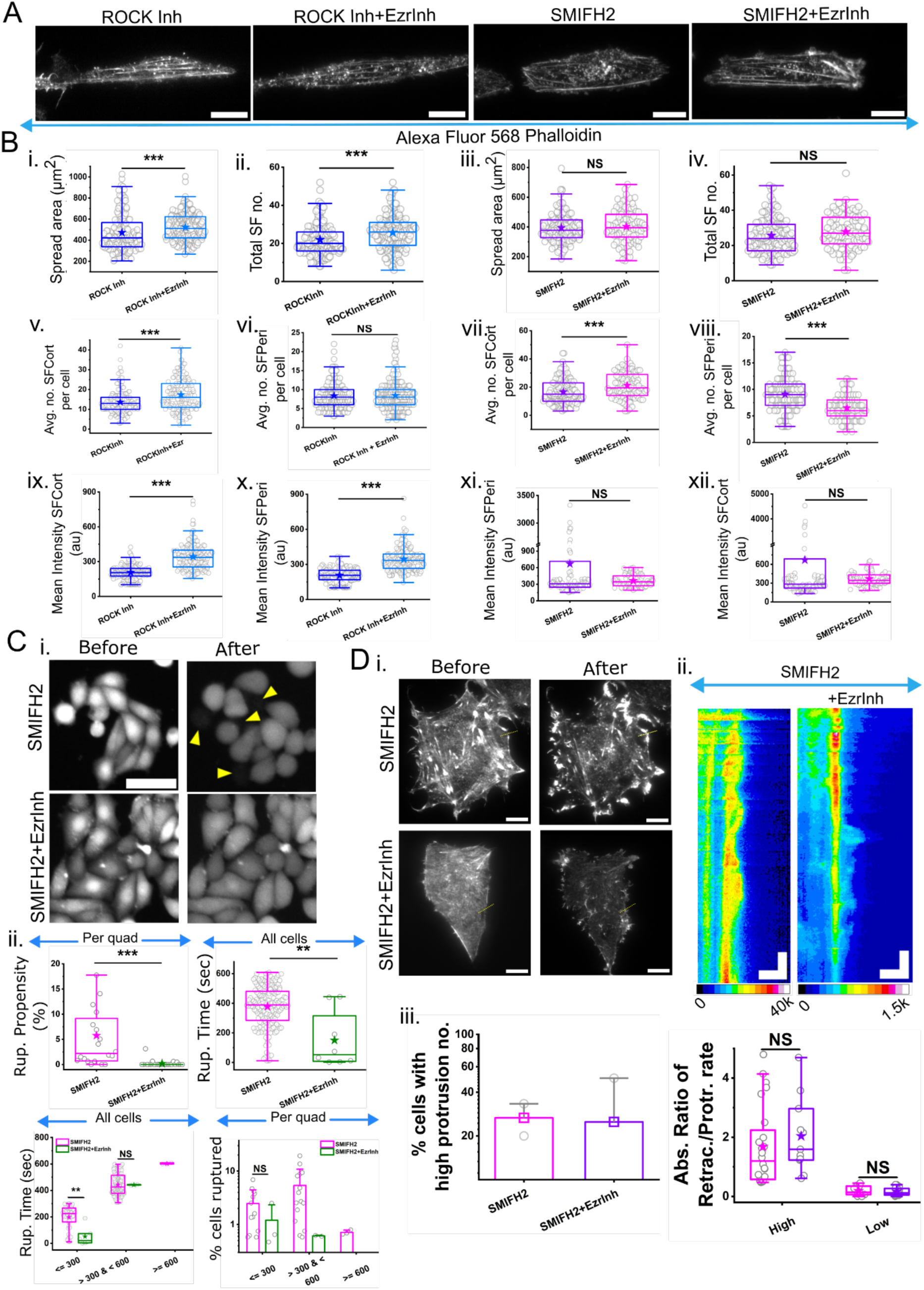
Stress fibre generation upon ezrin inhibition is majorly formin-mediated. A. Representative TIRF images of F-actin labelled with Alexa Fluor 568 Phalloidin ROCKInh (10 µM, 1 hr) or SMIFH2 (10 µM, 1 hr) pre-treated cells on EzrInh treatment. B. i-iv. Comparison of total number of stress fibres and spread area on EzrInh treatment. v-viii. Comparison of number of SFPeri and SFCort on ezrin inhibition. ix-xii. Comparison of mean intensities of SFPeri and SFCort. N= 3 independent experiments. n_ROCKInh_ = 157 cells, n_ROCKInh+EzrInh_ = 172 cells, n_SMIFH2_= 133 cells, n_SMIFH2+EzrInh_ = 94 cells. Scale: 10 µm. Statistical significance was determined by Mann Whitney U-test where *p < 0.05, **p < 0.01, ***p < 0.001 and ns p > 0.05 with Bonferroni correction. C. Rupture assay of CHO cells upon hypo-osmotic shock on EzrInh treatment in SMIFH2 pre-treated cells. i. Representative epi-fluorescence images of Calcein AM loaded cells before and after hypo-osmotic shock. ii. Comparison of rupture propensity (quadrant-wise), rupture time (all cells) and rupture time and percentage of cells rupturing across three time-scales. N= 3 independent experiments, No. of quadrants: Q_SMIFH2_ = 20, Q_SMIFH2+EzrInh_ = 24; Total number of cells n1, Total no. of ruptures: nrup _SMIFH2_ = 171, nrup _SMIFH2+EzrInh_ = 8. D. Representative TIRF images of SMIFH2 pre-treated CHO cells on EzrInh treatment, transiently expressing LifeAct mCherry, before and after hypo-osmotic shock. Scale: 10 µm. ii. Kymograph generated across line ROI (yellow line ROI in ii) representing evolution of protrusions (white arrows) and corresponding retractions (yellow arrows). X-axis represents distance across ROI (1 pixel= 65 nm) while time on Y-axis (1 pixel= 2 sec). Scale: 2 µm (X-axis), 60 sec (Y-axis). iii. Quantification of percentage of cells showing high protrusion numbers (>8 protrusions per cell). Also shown comparison of absolute ratio of rate of retraction and rate of protrusion. Error bars represent standard deviation. N= 2 independent experiments, n _SMIFH2_ = 5 cells, n_SMIFH2+EzrInh_ = 7 cells. Statistical significance was determined by Mann Whitney U-test where *p < 0.05, **p < 0.01, ***p < 0.001 and ns p > 0.05.

However, in SMIFH2 pre-treated cells, inhibiting ezrin neither enhanced spread nor total number of SFs (**Fig. 4B v,vi**). While the cortical SFs still increased the peripheral SFs showed a marked decrease. The mean intensities of SFs were enhanced on ezrin-inhibition in ROCK inhibited cells (**Fig. 4B ix, x**) but remained non-significant in SMIFH2 pretreated cells (**Fig. 4B xi, xii**). Our findings suggest that EzrInh mediated cortical stress fibre generation might involve both ROCK and formins however the balance of peripheral and cortical SFs also affecting cell spread area required formins.

If ezrin’s mechanoprotective role was regulated by formin-mediated balance of cortical and peripheral stress fibres^17,26^, we subjected SMIFH2 pretreated cells to hypo-osmotic shock. Notably, SMIFH2 treatment alone renders cells more susceptible to rupture (**Fig. 4C i**). However, when EzrInh is added to SMIFH2-pretreated cells, no additional increase in rupture is observed; instead, EzrInh appears to rescue cells from SMIFH2-induced vulnerability (**Fig. 4C i, ii**). This suggests that on ezrin inhibition resourcing of freed-up cortical actin to make cortical SFs and maintain peripheral SFs is mainly managed by formins. In the absence of formins actin at the cell edge on ezrin-inhibition becomes more homogeneous than contractile. Checking the retraction-to-protrusion rate ratio (**Fig. 4D i, ii**) revealed that neither of number of protrusions nor the ratio of rates was distinctly different in these two cases (**Fig. 4D iii**). Thus the rescuing action of formin inhibition was not be re-establishing contractility but stopping the increase in stress fibres on ezrin inhibition.

Together, this section could identify a major role played by formin in the cell’s reaction to ezrin inhibition. We bring out a new angle to the otherwise mechano-protective molecule formin. In ezrin inhibited conditions formins action of creating SFs from cortical actin creates inhomogeneities that makes the cell vulnerable to ripture during hypo-shock.

## Discussion

Our findings first demonstrate ezrin’s role in mechano-protection against hypo-osmotic shock in CHO cells. Inhibiting ezrin by its small molecule inhibitor disrupts actin-membrane interactions (witnessed as reduced membrane bound p-ezrin (**Fig. 1A**)) not only overall at the basal membrane but also at cell edges. This observation aligns with expectations, given ezrin’s known role in coupling the actin cytoskeleton to the plasma membrane. However, we show that inhibiting ezrin increases rupture propensity in the population significantly. While it is known that ERM (Ezrin Radixin Moesin) family of proteins have a role in cancer progression^27^ and also specifically ezrin inhibitors reduce invasiveness of osteosarcoma^28^, our work underscores that not only does ezrin inhibitor have the ability to delay motility (**Fig. S7F**) but also compromise cellular integrity.

We show that the mechanism is not a general increasing of membrane tension or a removal of actin from under the membrane. Rather, we report a change in the distribution of membrane tension. Similarly for actin, while the base saw an increase in SFs due to a cellular reaction to ezrin inhibition, the cortical actin at the midplane saw a weaker contractility. We captured the effect of hypo-shock on the tension reporter FliptR at the midplane of cells in a time dependent manner – a first report of time-dependence, to the best of our knowledge. However unlike IRM, FliptR FLIM is more phototoxic – hence could not be conducted on same cells through hypo-shock. Both IRM and FliptR could report that ezrin inhibition did not enhance the tension rise on hypo-shock. IRM also captured the edge to inner tension ratio at the basal plane - changing more in the first 10 minutes in ezrin inhibited cells. This is also evident from the tension maps of cells followed in time through hypo-shock (**Fig. 2C**). Following up regions that protrude out after hypo-shock, the actin dynamics revealed slower relative retraction of protrusions than control cells. The increase in number of protrusions was not significant but we found a greater percentage of cells in different experimental sets having >8 protrusions on hypo-shock. Thus, the reduction of contractility captured by myosin IIA FRAP experiments at the cell’s midplane was also reflected in the protrusion dynamics. While this recreates an image of a weaker actin network we also demonstrate a greater number of SFs on ezrin inhibition.

The increase in SFs was found to be concomitant with increased traction forces and reduced cell motility (**Fig. S7F**) – hence functionally relevant. We further revealed that the increase in SFs is majorly formin-dependent since formin’s pre-inhibition stops ezrin inhibition from increasing SFs. Although cortical SFs still increase – there is a decreases in the peripheral SFs and a lack of increase in spread area. Hence multiple parameters not only point to formin-dependent reaction to ezrin inhibition but also bring out the role of peripheral actin. This is in line with membrane dynamics showing that the peripheral membrane at the basal membrane too shows more relative enhancement in mechanical stress at 5 min post hypo-shock than control although the absolute values remain lower for ezrin-inhibited cells. The role of formin was validated by rupture experiments unambiguously demonstrating a reduction of rupture propensity (hypo-shock induced) on ezrin inhibition in formin-inhibited cells. While formins are known for their role in mechano-transduction^29^ – their coupling to ezrin-mediated mechno-protection is new. A careful investigation of different changes to the actin cytoskeleton by inhibition of ezrin could reveal the connection of formins in the mechano-sensitivity of cells. Stress fibres increase can protect cells by limiting their movement in this context – however, their over-formation in conjunction with a weaker cortex can make cells more susceptible to rupturing when stressed.

Throughout this work, we have explored the importance of tension heterogeneity or variations over basal average tension in understanding membrane resilience. Previous studies on liposome spreading^30^ have demonstrated that adhesion increases local mechanical stress, leading to rupture at contact sites. Our findings are in line with this showing in cells too enhanced cell spreading by EzInh correlate with enhanced ruptures while pre-treatment of cells with formin-inhibitor SMIFH2 converts actin to a more unform mesh preventing enhanced cell spreading and prevented ruptures.

## Material and Methods

### Cell culture

CHO cells were grown in growth media composed of Dulbecco’s Modified Eagle Media (DMEM, Gibco, USA) supplemented with 10 % foetal bovine serum (FBS, Gibco) and 1% Anti-anti (Gibco) at 37 °C, 5 % CO_2_ and 95% moisture. Cells were grown for at least 20 hours prior to using for experiments.

### Pharmacological interventions

To perturb ezrin, cells were pre-treated with Ezrin Inhibitor, NSC668394 (Sigma) at 40 µM concentration for 1 hour at 37°C. NSC668394 has been shown to inhibit ezrin effectively at 10 µM in osteosarcoma cells with other in vitro studies using concentrations up to 100 µM without any significant inhibition of PKC activity on ezrin^28^. Based on this, we used 40 µM for 1 hour in CHO cells to ensure robust inhibition without inducing cytotoxicity.

To disrupt actin stress fibers, a pretreatment with either formin inhibitor, SMIFH2 (Sigma) or ROCK inhibitor, Y27632 (Sigma) – both at concentration of 10 µM^31,32^ for 1 hour was given to cells.

### Fixation

CHO cells were fixed in 4 % PFA solution for 15 min. at 37 °C followed by quenching of autofluorescence from PFA by 0.1% glycine for 5 min. Next cells were treated with 0.1 M glycine (Sigma-Aldrich) for 5 minute followed by Triton-X (0.2%, 2 min) (Sigma-Aldrich). Blocking was done by incubating cells in 0.2% Gelatin (3 mi, overnight, 4°C) (Sigma-Aldrich). Cells were washed with 1X PBS at every step.

### F-actin labelling

F-actin labelling was done by incubating fixed cells with 1:200 dilution of Phalloidin Alexa Fluor 568 (Molecular Probes, Life Technologies) for 45 min. in dark. according to manufacturer’s instructions.

### Immunostaining

For p-ezr staining, fixed cells were incubated with primary antibody, Anti-Ezrin (phospho T567) (Abcam) (1:200, 3 hr), raised in rabbit, followed by a secondary antibody, Goat Anti-Rabbit IgG conjugated with Alexa Fluor 488 (1:1000, 2 hr).

For STED, cells are incubated with primary antibody as described above. Next cells are incubated with Abberior STAR Red–conjugated anti-rabbit secondary antibody (Abberior) (1:400, 1 hr). Finally, the samples were washed three times in 1× PBS and mounted onto clean cover-slips using MOWIOL 4-88 mounting medium (Sigma-Aldrich).

### Flow cytometry

Well spread CHO cells growing on 60 mm dishes are fixed with 4% PFA for 15 mins. And then scrapped gently with a cell scrapper. Scraped cells are collected in PBS and centrifuged at 1500 rpm for 5 min and then subjected to p-ezr antibody staining protocol described in previous section. Cells were finally washed with PBS and resuspended in 500 μl PBS then analysed by flowcytometry (BD FACSDiva 9.0). Secondary antibody tagged with Alexa Fluor 488 was excited with 488 nm laser and emission signal is collected with 527/32 band pass filter Same parameters such as PMT voltage, threshold and gating was used throughout all conditions. For every condition 10,000 cells were analysed.

### Transfection

For live cells visualization of F-actin, well spread CHO cells growing on glass cover-slips were transfected with pDEST/Life-Act-mCherry-N1(a gift from Rowin Shaw, Addgene, plasmid#40908). For visualization of myosin IIa, Lin10 micropatterned CHO cells were transfected with mCherry-MyosinIIA-C-18 (a gift from Michael Davidson, Addgene, plasmid#55105). Cells were seeded for at least 16 hr prior to transfection. All transfections were done using 1 µg of plasmid DNA by lipofection (Lipofectamine 3000, Life Technologies).

### Hypo-osmotic shock and Rupture analysis

Cells grown on glass cover-slips were washed with PBS and M1 media added to them. For hypo-osmotic shock, DMEM diluted in deionized water (1/20× for 95% shock) is used as the hypo-shock media. For control condition, 2.5 µM Calcein AM (Invitrogen) and Hoechst 33342(Invitrogen) at 0.5 µg/ml working concentrations are added to the cells along with the M1 media for 30 mins at 37°C. For drug treated condition, Calcein AM and Hoechst 33342 is added after 30 mins of pretreatment with drugs. The treated cells are taken for timelapse imaging.

First few frames are captured then existing media is changed and hypo-shock media is added to the cells. The time between discarding existing media and application of hypo-shock media is the adjustment factor. To calculate rupture propensity, cells are scanned for 10 mins after the hypo-osmotic shock and cells losing Calcein AM are counted as rupturing cells.

First a few frames were captured then existing media was changed and hypo-shock media was added to the cells. The time between discarding existing media and application of hypo-shock media was considered as the adjustment time. To calculate rupture propensity, cells were scanned for 10 min after the hypo-osmotic shock and cells losing Calcein AM were counted as rupturing cells.

Image analysis was performed using Image J/Fiji. To calculate total no. of cells/quadrant (Nt), the imaged field was divided into 4 quadrants (∼ 100 cells per quadrant) and the number of Calcein AM loaded cells with nuclei stained with Hoechst were counted for each quadrant. From epifluorescence images, number of cells which loses Calcein AM signal are marked as the rupturing cells (Nr). Rupture Propensity (Rp) per quadrant is calculated as: *Rp* = (*Nr*/ *Nt*) × 100 %.

Rupture time was calculated by scanning the timelapse images to mark the exact frame (Fr) where Calcein AM signal is lost for the rupturing cell. To account for the time between discarding existing media and application of hypo-shock media is the adjustment time, the rupture time (Rt) is calculated as:*Rt* = (*Fr* × 2 *sec*) + *Adjustment time*. Adjustment time was same for cells foor a given dish but the rupture time varied from cell to cell.

### TIRF Microscopy

TIRF imaging was done on an inverted microscope (IX81, Olympus Corporation, Japan) using a 100x TIRF objective with 1 pixel = 65 nm and a CMOS camera (ORCA-Flash 4.0, Hamamatsu Photonics, Japan) using lasers of 561 nm wavelengths and at penetration depth of 72 nm. For all live cell imaging cells were maintained at 37°C throughout by using an onstage as well as cage incubator (Okolab, Italy). Timelapse images were captured as per described in rupture analysis.

### P-ezrin Image Analysis

P-ezrin fluorescence intensity analysis was performed on background-subtracted images. A rectangular region around each cell was manually selected, and a smaller background ROI was chosen to estimate background intensity, which was subtracted from the image. The full cell area was then outlined using a freehand ROI to define the total cell mask. Within this mask, the mean P-ezrin intensity was calculated. To isolate the inner cortical region, the whole-cell mask was eroded by 30 pixels (corresponding to ∼2 µm), and the difference between the original and eroded masks was used to generate a narrow inner membrane ring. The median fluorescence intensity within the resulting ROI was then calculated for each image.

### FRAP

Lin 10 micropatterned CHO cells were transfected with mCherry-MyosinIIA-C-18 (a gift from Michael Davidson, Addgene, plasmid#55105). FRAP experiment was carried on as per described protocol^22^.

### Confocal FLIM live imaging of FliptR

FLIM imaging was performed using the Abberior Facility Line system integrated with an Olympus IX83 inverted microscope, equipped with a time-correlated single photon counting (TCSPC) module from PicoQuant. Excitation was carried out using a pulsed 488 nm diode laser, operated at a repetition rate of 40 MHz. Data acquisition and control were managed through PicoQuant’s SymPhoTime software. Images were acquired at a lateral resolution of 100 nm per pixel.

To probe membrane tension and lipid packing, cells were incubated with 1 µM FliptR in M1 medium for 40–50 minutes at 37 °C. After incubation, excess dye was removed by washing with fresh M1 medium. Imaging was conducted in Ibidi two-well glass-bottom dishes under live-cell conditions.

For hypo-osmotic shock, same setup and protocol was used. However, cells were grown on glass cover-slips mounted on custom-made glass-bottom dish with silicone gel A 95% hypo-osmotic shock was given, and cells were imaged at three time points: before the shock (baseline), 5 minutes after shock, and 20 minutes after shock. Imaging was conducted at the mid-cell plane with an acquisition pixel size of 200 nm.

### Lifetime Image Analysis

For analysis, the phasor-analysis-derived lifetime was utilized considering only pixels with photon count greater than 100. A Gaussian filter was first applied to suppress background variations, and the result was subtracted from the original image to enhance edge features. The processed image was normalized and binarized using a fixed intensity threshold, followed by removal of small isolated regions below a defined area cutoff. A cell boundary was manually traced using a freehand ROI on the intensity image to remove non-cell regions. A cortical mask approximately 2 pixels wide (∼200 nm) was generated by morphologically eroding the segmented cell edge. This mask was then applied to the corresponding lifetime image to extract lifetime values specifically along the cell cortex. The median fluorescence lifetime within the resulting cortical ROI was then calculated for each image.

### IRM imaging and analysis

Cell imaging was performed using a Nikon Eclipse Ti-E motorized inverted microscope (Nikon, Japan), equipped with adjustable field and aperture diaphragms. A 60× Plan Apo water immersion objective (NA 1.22) and an additional 1.5× external magnification were used. The microscope is equipped with an on-stage incubator maintained at 37°C (Tokai Hit, Japan). Images were acquired with an sCMOS camera (ORCA Flash 4.0, Hamamatsu, Japan).

Illumination was provided by a 130 W mercury arc lamp, utilizing an interference filter (546 ± 12 nm) and a 50:50 beam splitter. For Interference Reflection Microscopy (IRM), time-lapse movies consisting of 2048 frames were captured at 20 frames per second, with an exposure time of 50 ms per frame.

IRM image analysis to extract membrane fluctuation tension was carried out in three major steps. First, a calibration factor was determined to convert intensity values into relative height measurements. Pixels corresponding to the first branch region (FBR) of the interference pattern were identified^33,34^. 12×12 pixel regions were grouped as FBRs used for further analysis. Note that 1 pixel = 74 nm. Next, a time series of height values was obtained for each pixel, and the amplitude of fluctuations—both temporal (SD_time_) and spatial (SD_space_)—was calculated within the FBRs. Finally, for each FBR, the power spectral density (PSD) was computed and fitted using a modified Helfrich–Canham model^35^ to extract the fluctuation tension and other membrane mechanical parameters. FBRs located within 10 pixels of cell boundary (drawn manually) were taken as edge-FBRs while others were considered as inner-FBRs.

For the experiment, CHO cells were serum-starved in M1 medium for 1 hour. In the case of Ezrin-inhibited cells, the serum-starved M1 medium was supplemented with 40 μM Ezrin Inhibitor. Before the hypoosmotic shock, an IRM movie consisting of 2048 frames was recorded. Following a 95% hypoosmotic shock, additional IRM movies (2048 frames each) were acquired at 2, 5, 10, and 20 min post-shock.

### Calculation of protrusions

From timelapse images of cells during hypo-osmotic shock, regions showing protrusion and retraction over time are isolated. Line ROIs were drawn perpendicular to the protruding edge and kymograph generated for such lines. The X axis denotes distance covered and Y axis denotes time elapsed. In the kymograph, using angle tool in ImageJ/Fiji, ROIs were drawn over the protruding and retracting edges and the respective angles (*θ*) noted. From these angles, rate of protrusion or retraction is calculated as 1/tan(*θ*).

### Stress fibre analysis

Image analysis was performed using Image J/Fiji. To calculate mean length and number of stress fibres, the analysis was performed in two parts-smoothening and background subtraction for better visualization of stress fibres and in the next part, drawing the ROIs over the stress fibres and measuring the number and length of each stress fibre. Also, parameters like spread area and total intensity are calculated for each cell.

### Micropatterning, hydrogel preparation and Traction force microscopy

Etched glass cover-slips were exposed to deep UV in the UV-Ozone cleaner (Jelight Company, USA) for 5 min to clean the surface of cover-slips. These cover-slips were next incubated with 0.2 mg/ml PLL-g-PEG (SuSoS, Switzerland) (prepared in 10 mM HEPES (Sigma), pH 8.5) for 4 h. Photo-masks (JD Photo Data, UK) were first cleaned in the UV-Ozone cleaner for 5 min and then used for patterning the PLL-g-PEG coated cover-slips for 5 min. PLL-g-PEG coated cover-slips were then placed on the photomask with coated the coated surface facing towards the photomask. A drop of water was added to the cover-slip before placing on photomask to aid in adherence. Patterned cover-slips were removed from photomask by floating them off using water. Cells were seeded on micropatterned cover-slips after washing with PBS.

For traction force microscopy, polyacrylamide hydrogel was prepared with final working concentration of 7.25% acrylamide (BioRad, Hercules, CA) and 0.16% Bis-acrylamide (BioRad). A gel solution was prepared by mixing 0.01% of 100 nm fluorescent beads (Molecular Probes, Life Technologies), 0.5% ammonium persulfate (Calbiochem), 0.05% tetramethylethylenediamine (Sigma) with the acrylamide and Bis-acrylamide mixture. 7 μl of the final polymerizing gel mixture were sandwiched between micropatterned circular cover-slip (10 mm diameter) and treated square cover-slips (22 × 22 mm) and kept in an inverted position such that the fluorescent beads would be localized at the surface of the hydrogel. After polymerization of hydrogel, micropatterned circular cover-slip was removed carefully and micropatterned hydrogel containing cover-slips was mounted on custom-made glass-bottom dish with silicone gel followed by cell seeding.

Traction force imaging was done by 60 × 1.49 NA objective. Bright field image and corresponding fluorescence image of beads embedded beneath the cell was captured. Next cells were removed by trypsinizing them and image of beads in relaxed condition was taken. From the displacement of beads, the traction force is calculated.

### Stimulated Emission Depletion (STED) Imaging

STED microscopy was performed using the Abberior Facility Line system integrated with an Olympus IX83 inverted microscope. Alignment between the confocal and STED point spread functions was verified using multicolor fluorescent calibration beads. Images were acquired at a lateral resolution of 20 nm per pixel, focusing on two optical planes: the basal and mid-cell levels (∼3 um above the base). A pulsed 775 nm laser was used for depletion of red and far-red fluorophores. F-actin was stained with Alexa Fluor 568–conjugated phalloidin. Phosphorylated ezrin (P-ezrin) was detected using a primary antibody and visualized with an Abberior STAR Red–conjugated anti-rabbit secondary antibody.

### Statistical Analysis

Statistical significance of all measurements was determined by Mann Whitney U test. For quantification of stress fibres, Bonferroni correction was also done along with Mann Whitney U test.

## Acknowledgement

B.S. acknowledges support from Wellcome Trust/DBT India Alliance fellowship (grant number IA/I/13/1/500885) for the TIRF set up. A.B. thanks UGC for providing her fellowship. The authors are also thankful to the Builder Imaging facility (BT/INF/22/SP45383/2022) for super-resolution STED imaging and Central Imaging Facility, DBS, IISER Kolkata for FRAP and Flow cytometry.

## Author Contributions

B.S., designed research, acquired funding; B.S., A.B., and R.K. set up methodology; A. B., R.K. H.T.N., J.D. performed research and analysed data; and A.B. and B.S. wrote the paper; everyone edited the paper.

## Data and resource availability

All relevant data and details of resources can be found within the article and its supplementary information.

Relevant codes for analysis are available at : https://github.com/BidishaSinha/Ezrin-Paper-2025

