## Supplementary Text for "Inhibiting ezrin triggers formin-mediated actin remodelling reducing cellular mechano-protection"

List of Figures:

**Figure S1**. **Inhibiting ezrin reduces p-ezrin**.
**Figure S2. Assaying hypo-osmotic shock-based rupture assay**.
**Figure S3. Distributions of rupture times.**
**Figure S4. Membrane parameters during hypo-shock.**
**Figure S5. Kymographs and absolute ratio of retraction to protrusion rates.**
**Figure S6. Montage of micropatterned cells with F-actin labelled.**
**Figure S7**. **Inhibition of ezrin reduces cellular motility.**
**Figure S1. Effect of ROCK inhibition and formin inhibition on F-actin.**
**Figure S*9*. SF characteristics on ROCK/Formin inhibition.**

**Table S1: List of statistical parameters of relevant main figures**
**Table S2: List of statistical parameters of relevant supplementary figure**

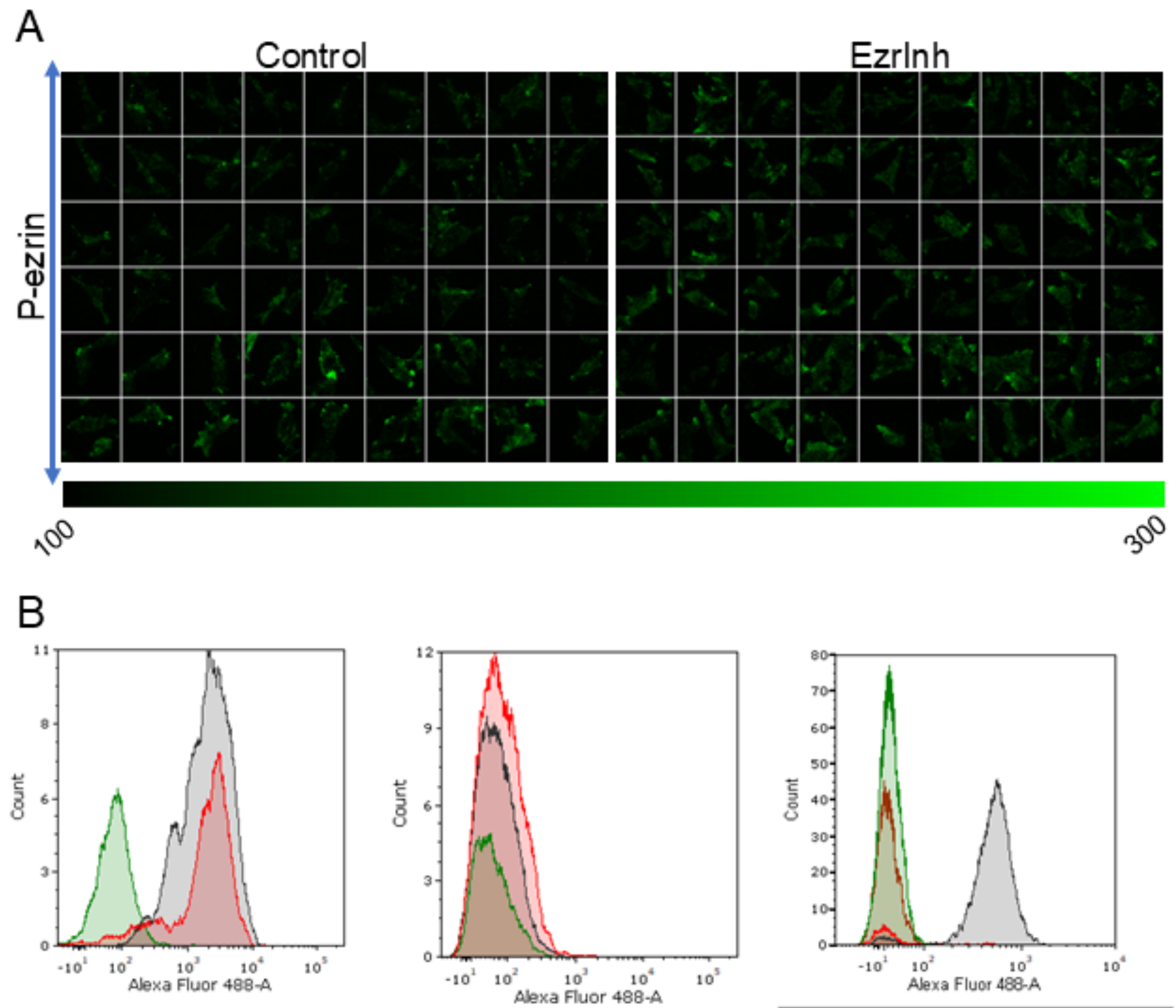

| Overlay color | Filename | # of Events | Median | Arithmetic Mean |
|---|---|---|---|---|
| | Ezr_SecA488A594.fcs | 754 | 64.80 | 77.98 |
| | Ezr_60min.fcs | 1112 | 1773.09 | 2669.21 |
| | Ezr_00min.fcs | 2205 | 1624.86 | 2360.98 |

| Overlay color | Filename | # of Events | Median | Arithmetic Mean |
|---|---|---|---|---|
| | Specimen_001_Ctl.fcs | 1584 | 52.65 | 82.17 |
| | Specimen_001_SecA488.fcs | 744 | 40.50 | 234.83 |
| | Specimen_001_60min.fcs | 2154 | 63.18 | 131.57 |

| Overlay color | Filename | # of Events | Median | Arithmetic Mean |
|---|---|---|---|---|
| | CHO Unst2.fcs | 3717 | 14.11 | 16.96 |
| | CHO 2Ct2.fcs | 6294 | 19.92 | 27.05 |
| | CHO Expt ctl2.fcs | 6177 | 500.49 | 521.90 |
| | CHO EI2.fcs | 602 | 19.09 | 262.42 |

**Figure *S1*. Inhibiting ezrin reduces p-ezrin**. A. Montage of TIRF images of CHO cells grown on glass coverslips and P-ezr stained with Rb mAb to Phospho-ezrin (T257) followed by Goat pAb to Rb IgG conjugated with Alexa Fluor 488. Cells are pretreated with EzrInh (40 μM, 1 hr) prior to fixation with PFA (4%, 15 min) followed by antibody staining. N= 2 independent experiments, $n_{control}$ = 40 cells, $n_{EzrInh}$ = 28 cells. B. Total P-ezr levels in EzrInh CHO cells determined by Flow cytometery. N=3 independent experiments.

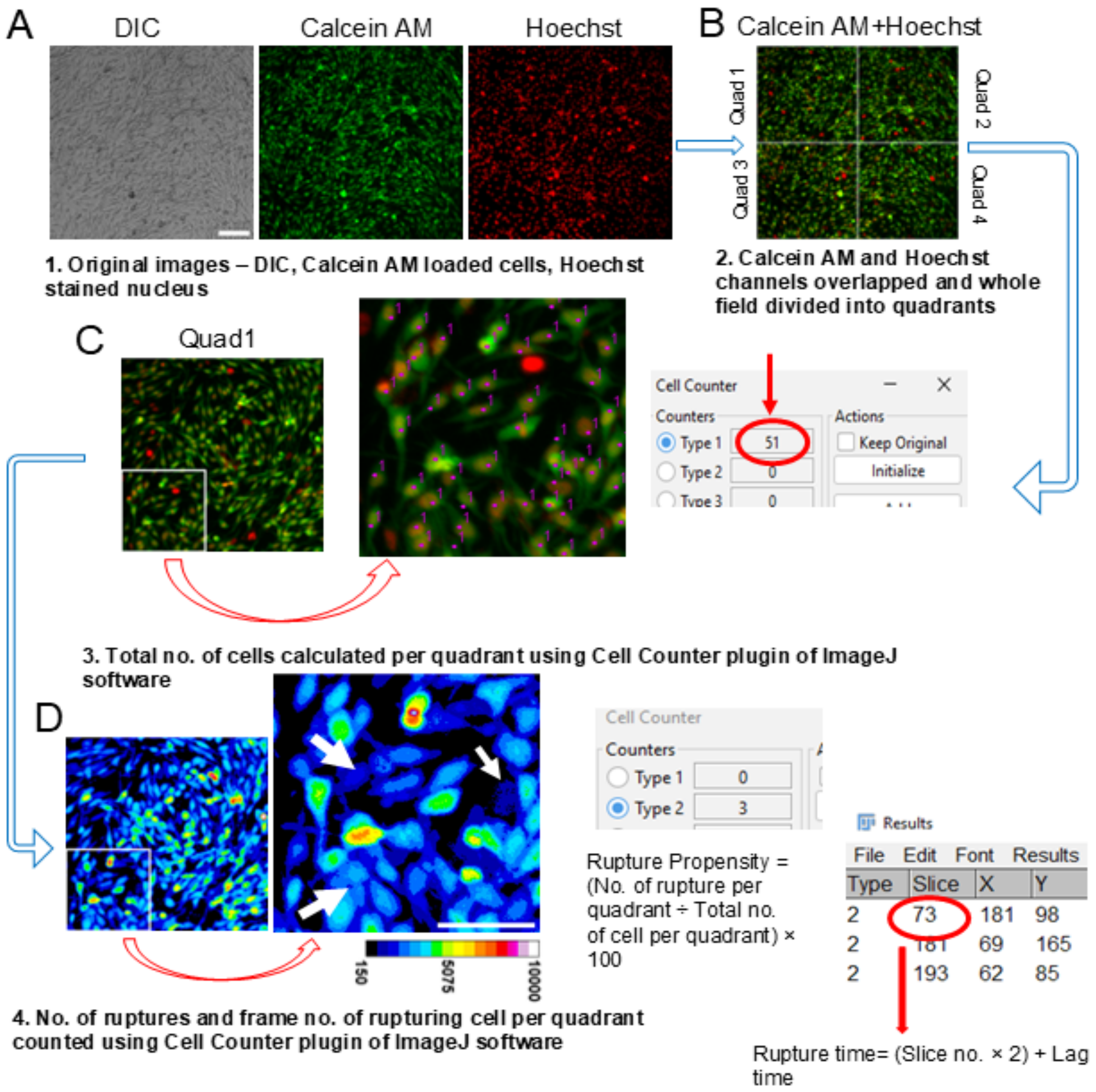

**Figure *S2*. Assaying hypo-osmotic shock-based rupture assay**. A. Epifluorescent images of Calcein AM loaded CHO cells is captured on an inverted microscope (IX81, Olympus Corporation, Japan) using 10×Plan Apo objective (NA 1.0003) and a CMOS camera (ORCA-Flash 4.0, Hamamatsu Photonics, Japan) at 0.5 frames/s for 10 min with 1 pixel=650 nm. Also, cell nucleus is stained with Hoechst stain. B. Calcein AM and Hoechst images are overlapped and the field is divided into four quadrants. C. Now each quadrant is run through Image J/Fiji plugin, “Cell Counter”. From the plugin, total number of cells per quadrant is calculated. D. For counting the number of ruptures and rupture time, timelapse images of each quadrant is run through the mentioned plugin multiple times and rupturing cells and the frame of rupture is recorded for further analysis.

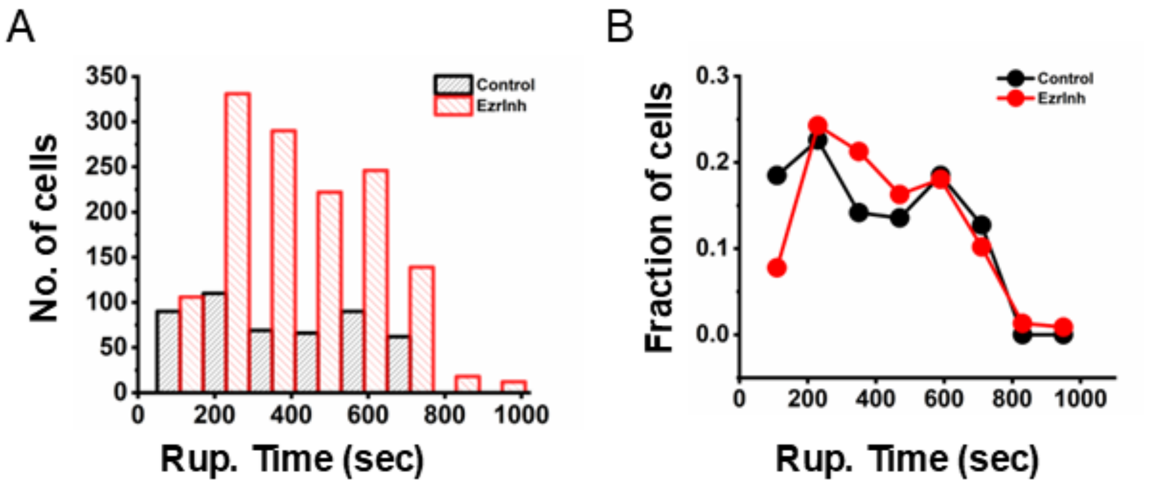

**Figure *S3*. Distributions of rupture times.** A. Comparison of distribution of rupture times of ruptured cells upon EzrInh treatment. B. Cells show a bimodal distribution indicating that ezrin inhibition affects rupture time across a range of time-scales. Bin size: 120. Total no. of ruptures: $nrup_{Control}$ = 487, $nrup_{EzrInh}$ = 1364.

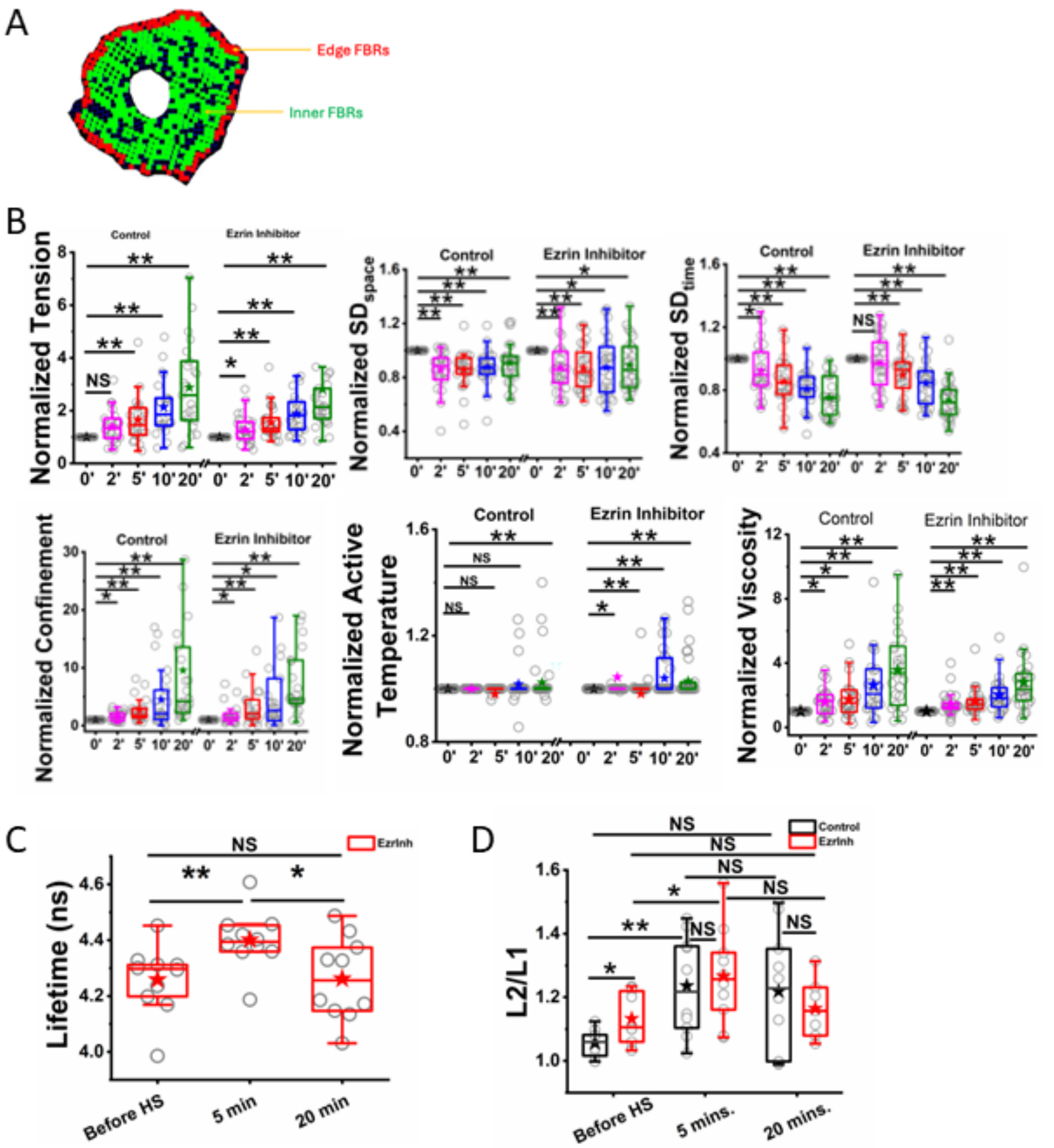

**Figure *S4*. Membrane parameters during hypo-shock.** (A) Representative image of a cell analyzed for IRM with edge FBRs (square ROIs of dimension 12x12 pixels or 864x864 nm2) depicted in red and inner FBRs depicted in green for tension ratio calculation. If any pixel of an FBR was located within 10 pixels (or 720 nm) of the cell boundary, the FBR was considered as an edge FBR. (B) pooling single cells followed in time, the trajectory of each cell was normalized to its before-shock time point. Normalized values depicting fold change in all associated mechanical parameters of the PSD fitting. N=4 independent experiments, $n_{Control}$ = 23 cells, $n_{EzrInh}$ = 24 cells with same cells followed over time. Statistical significance was determined by Mann Whitney U-test where $*p < 0.05$, $**p < 0.001$, and ns $p > 0.05$. (C, D) Evolution of lifetime of FliptR (C) and in ratio of mid-to-base lifetime of FliptR (L2/L1). (D)

with time after hypo-shock. N=1 repeat. $n_{Control}$ = 10 cells, $n_{EzrInh}$ = 10 cells. Statistical significance was determined by Mann Whitney U-test where *$p < 0.05$, **$p < 0.01$, ***$p < 0.001$ and ns $p > 0.05$.

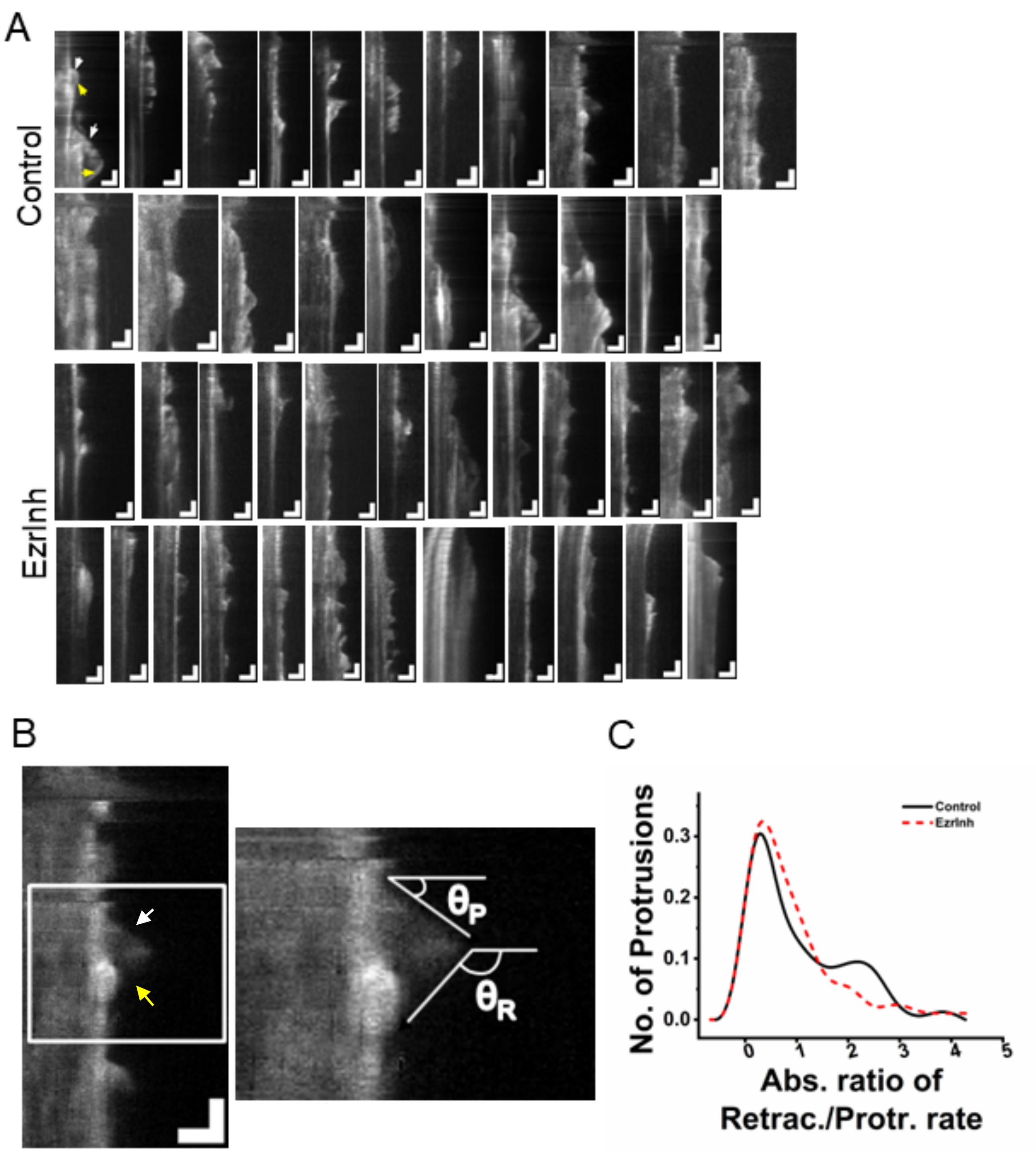

**Figure *S5*. Kymographs and absolute ratio of retraction to protrusion rates.** A. Montage of kymographs used to quantify protrusions and retractions upon hypo-osmotic shock in ezrin inhibited cells. X-axis represents distance across ROI (1 pixel= 65 nm) while time on Y-axis (1 pixel= 2 sec). Scale: 2 μm (X-axis), 60 sec (Y-axis). B. (*right*) Representative kymograph showing protrusions (*white arrow*) and retractions (*yellow arrow*). Protrusion angle ($\theta_P$) and retraction angle ($\theta_R$) calculated with "Angle tool" in Image J/Fiji (zoomed-in portion, *left*). C. Comparison of normalized distribution of absolute ratio of rate of retraction and rate of protrusion on EzrInh treatment.

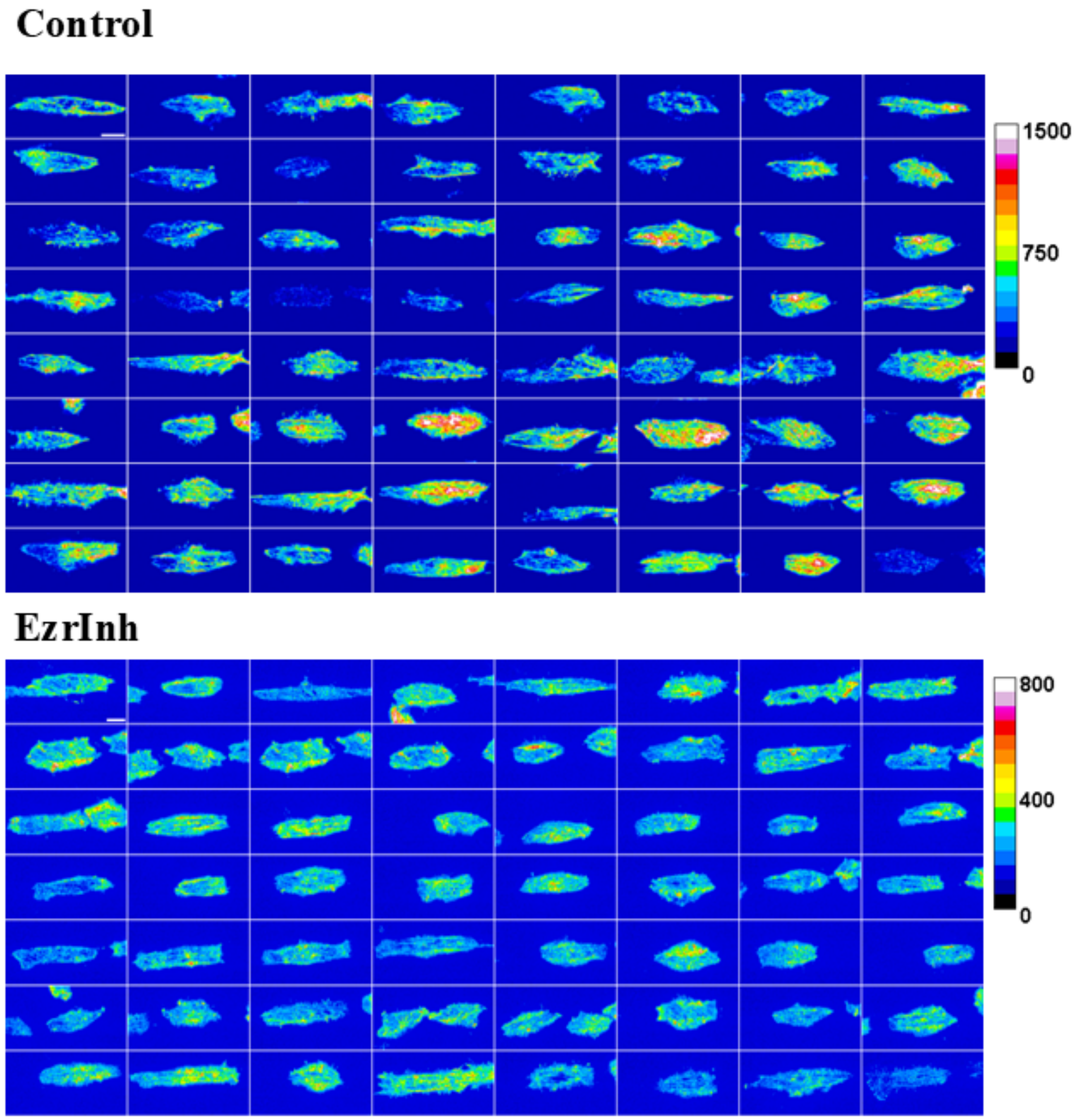

**Figure *S6.* Montage of micropatterned cells with F-actin labelled.** TIRF images of Lin 10 micro-patterned CHO cells with F-actin labelled with Alexa Fluor 568 Phalloidin. EzrInh treatment increases stress fibres at basal plane. F-sctin intensity is denoted by associated colorbar. N= 3 independent experiments, $n_{Control}$=169 cells, $n_{EzrInh}$=175 cells. Scale: 10 µm.

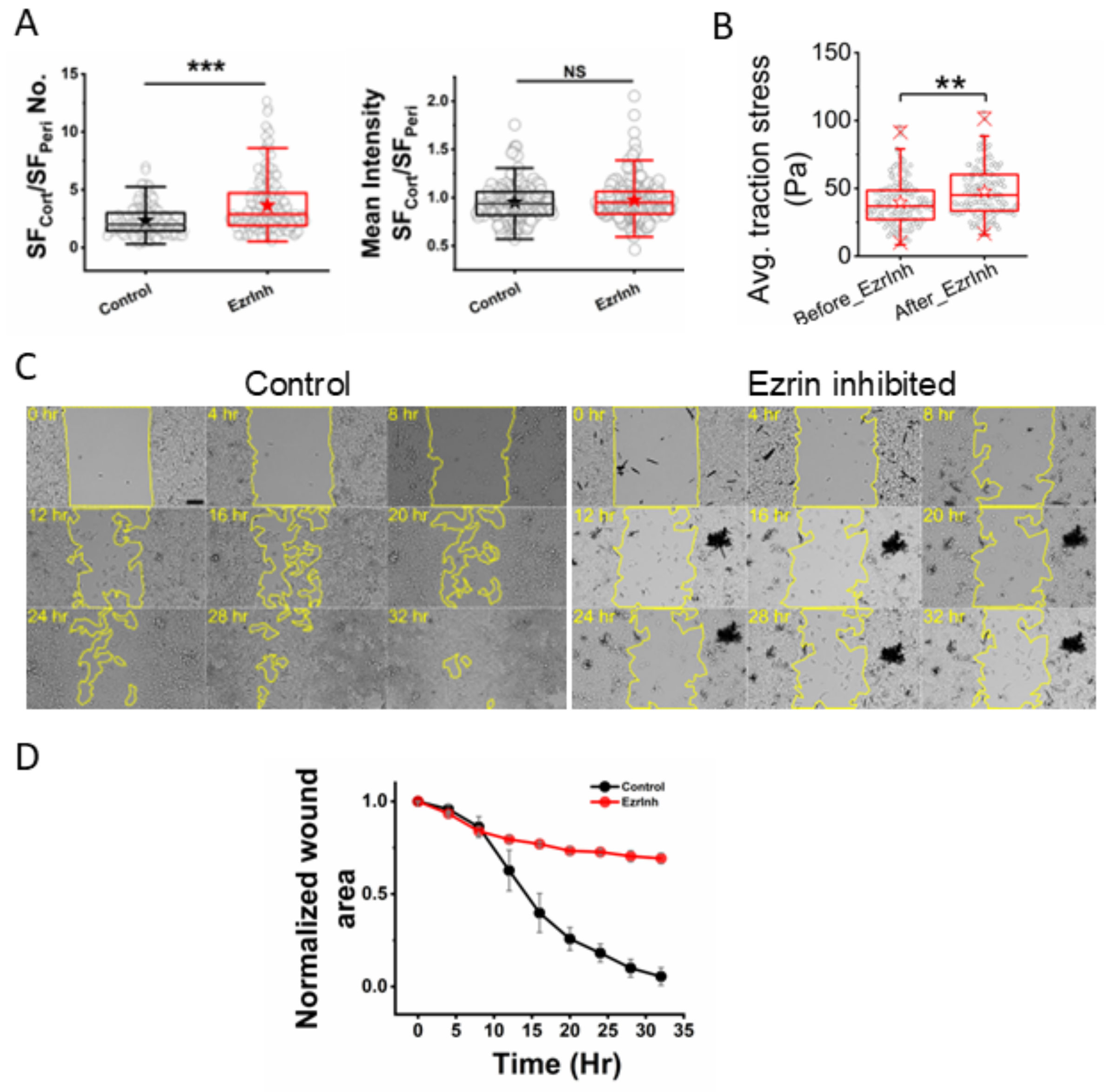

**Figure** ***S7*****. Inhibition of ezrin reduces cellular motility.** A. Comparison of ratio of number and mean intensity of SFCort and SFPeri on ezrin inhibition. Statistical significance was determined by Mann Whitney U-test where $*p < 0.05$, $**p < 0.01$, $***p < 0.001$ and ns $p > 0.05$, with Bonferroni correction. B. Comparison of average traction force on EzrInh treatment. C. Representative images of wound healing on ezrin inhibition. D. Comparison of wound closure over time. N= 1 independent repeat. 3 wound area analyzed per condition. Wound healing assay has been done by growing CHO cells in ibidi 35 µm cell culture dish with 2 well inserts. The inserts were carefully removed after cells were well spread (wound area) and fresh media added. The cells were followed for 32 hours with 4 hour intervals under an inverted microscope (IX81, Olympus Corporation, Japan) using 10×Plan Apo objective (NA 1.0003) and a CMOS camera (ORCA-Flash 4.0, Hamamatsu Photonics, Japan) with 1 pixel= 650 nm.

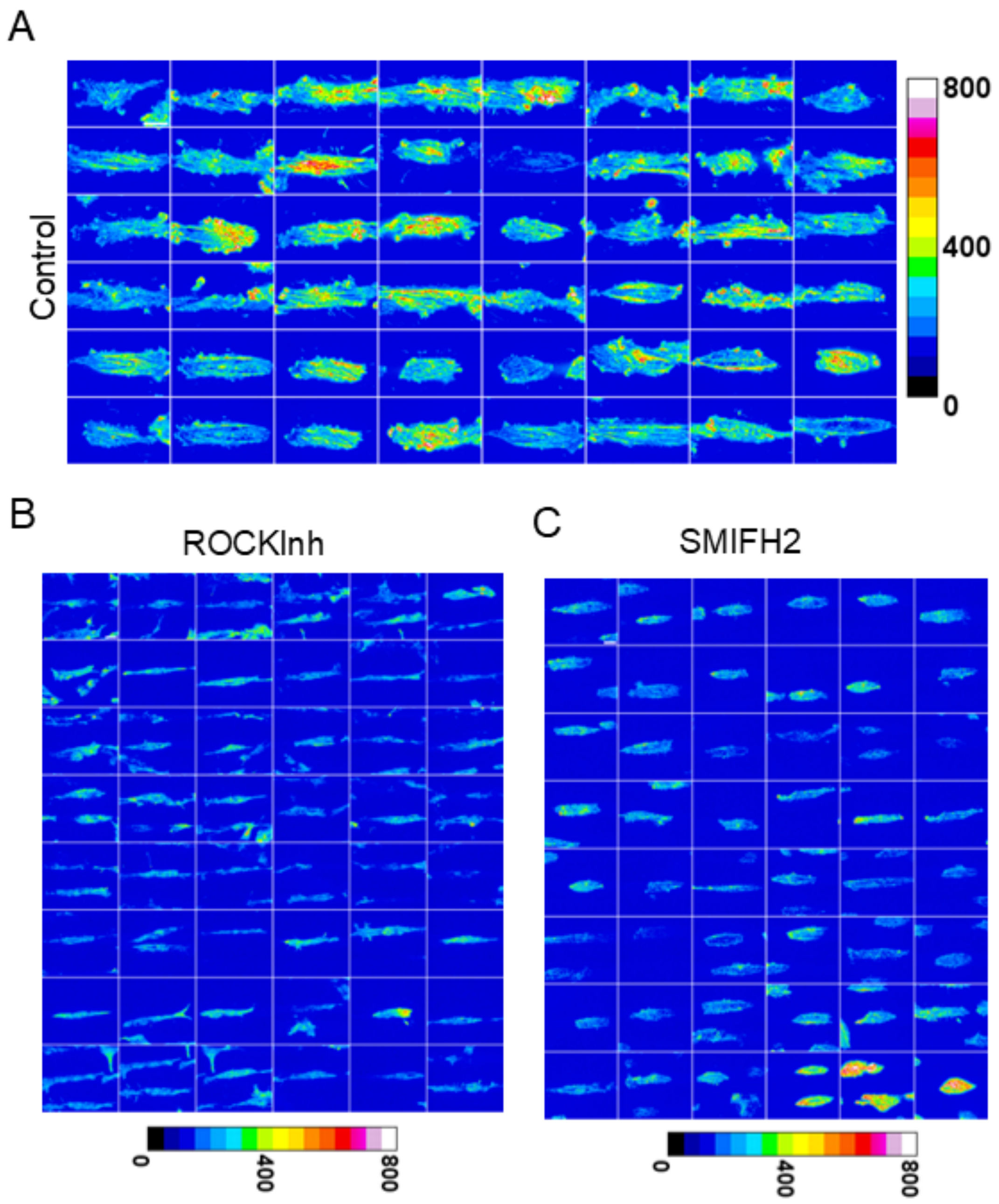

**Figure *S2*. Effect of ROCK inhibition and formin inhibition on F-actin.** Montage of Lin 10 micro-patterned CHO cells pretreated with ROCK Inh/ SMIFH2 (10 µM, 2 hr) prior to fixation and labelling of stress fibres with Alexa Fluor 568 Phalloidin. 16-color LUT applied to images and F-sctin intensity is denoted by associated colorbar. N= 3 independent repeats; $n_{Control}$=169 cells, $n_{ROCKInh}$= 157 cells, $n_{SMIFH2}$= 133 cells. Scale: 10 µm.

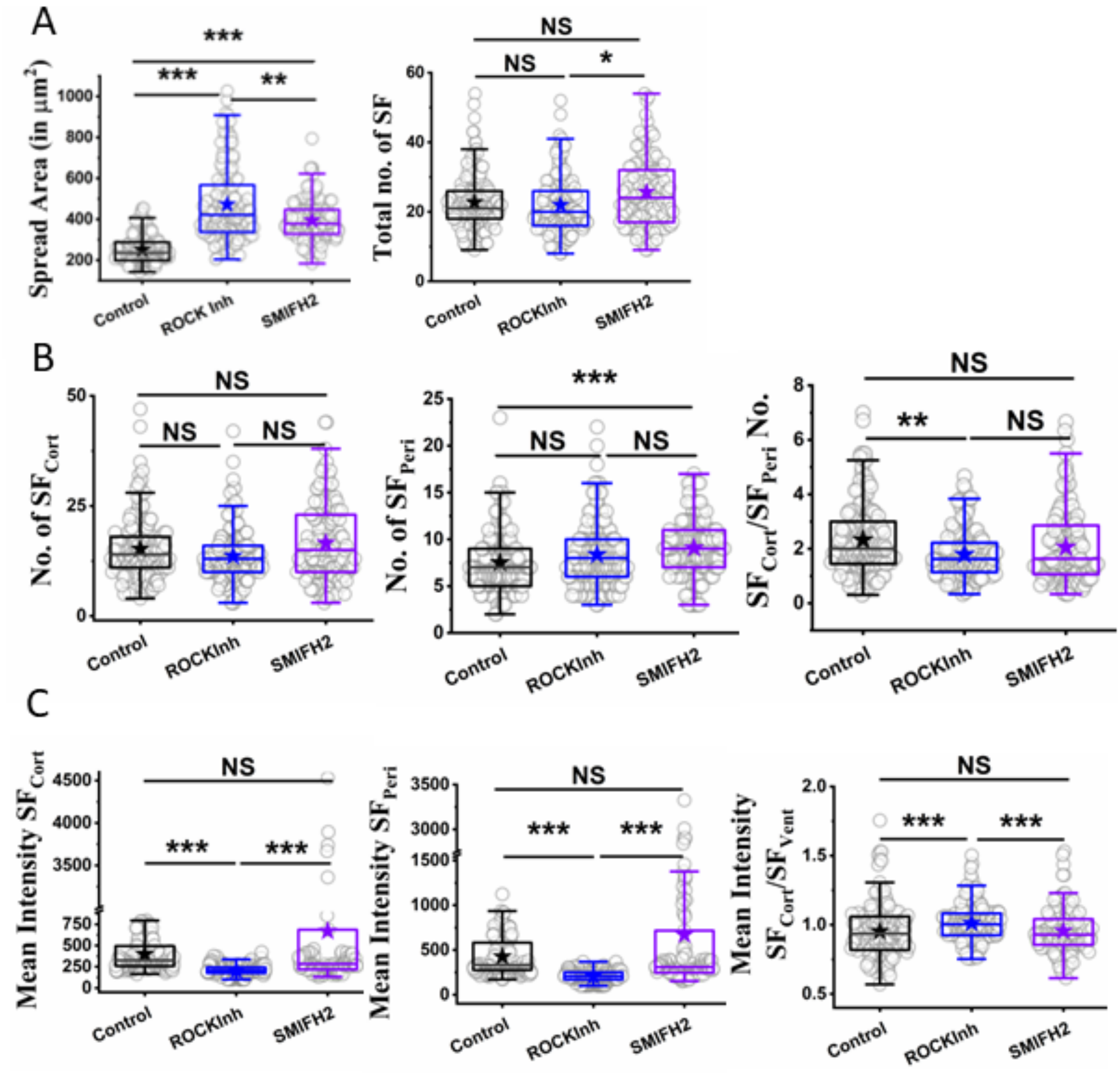

**Figure S9. SF characteristics on ROCK/Formin inhibition.** Comparison of stress fiber parameters on ROCKInh (10 μM, 1 hr) or SMIFH2 (10 μM, 1 hr) treatment. N= 3 independent repeats; $n_{Control}$=169 cells, $n_{ROCKInh}$= 157 cells, $n_{SMIFH2}$= 133 cells. Statistical significance was determined by Mann Whitney U-test where *p < 0.05, **p < 0.01, ***p < 0.001 and ns p > 0.05, with Bonferroni correction.

Table S1

| Figure 1E | | | | | | | | | | |
|---|---|---|---|---|---|---|---|---|---|---|
| **Parameters** | **Conditions** | **Set** | **Dish** | **Quadrant** | **ntotal** | **nrup** | **Median** | **p-values** | **Trend** | **Fractional change** |
| Rupture propensity | Control | 9 | 16 | 64 | 18603 | 487 | 1.08 | | | |
| | EzrInh | 9 | 23 | 92 | 30557 | 1364 | 2.05 | 0.00 | Increases | 0.90 |
| Rupture Time | Control | 9 | 16 | 64 | 18603 | 487 | 361.00 | | | |
| | EzrInh | 9 | 23 | 92 | 30557 | 1364 | 387.00 | 0.00 | Increases | 0.072 |
| **Figure 1F** | | | | | | | | | | |
| **Parameters** | **Conditions** | **Set** | **Dish** | **Quadrant** | **ntotal** | **nrup** | **Median** | **p-values** | **Trend** | **Fractional change** |
| Rupture time categorised (in sec) | | | | | | | | | | |
| <= 300 | Control | 9 | 16 | 64 | 16055 | 214 | 186.00 | | | |
| <= 300 | EzrInh | 9 | 23 | 92 | 30557 | 467 | 208.00 | <0.0001 | Increases | 0.12 |
| > 300 & < 600 | Control | 9 | 16 | 64 | 18603 | 173 | 469.00 | | | |
| > 300 & < 600 | EzrInh | 9 | 23 | 92 | 29766 | 624 | 440.50 | 0.02 | Decreases | -0.06 |
| >= 600 | Control | 9 | 16 | 64 | 15977 | 100 | 658.00 | | | |
| >= 600 | EzrInh | 9 | 23 | 92 | 30557 | 273 | 670.00 | 0.07 | NS | |
| % cells ruptured (per quad) | | | | | | | | | | |
| <= 300 | Control | 9 | 11 | 33 | 1074 | 214 | 0.76 | | | |
| <= 300 | EzrInh | 9 | 21 | 58 | 20219 | 467 | 1.14 | 0.14 | NS | |
| > 300 & < 600 | Control | 9 | 14 | 38 | 12047 | 173 | 0.73 | | | |

| | | | | | | | | | | |
|---|---|---|---|---|---|---|---|---|---|---|
| > 300 & < 600 | EzrInh | 9 | 18 | 66 | 22113 | 624 | 1.23 | 0.03 | Increases | 0.67 |
| >= 600 | Control | 9 | 11 | 24 | 6961 | 100 | 0.62 | | | |
| >= 600 | EzrInh | 9 | 21 | 66 | 21555 | 273 | 0.87 | 0.48 | NS | |
| **Figure 2A ii.** | | | | | | | | | | |
| **Parameters** | **Conditions** | **Set** | **Mean** | **SD** | **ntotal** | **Median** | **SEM** | **p-values** | **Trend** | **Fractional change** |
| tau | Control | 2 | 3.83 | 1.14 | 40 | 0.18 | 3.78 | | | |
| | EzrInh | | 3.17 | 0.98 | 42 | 0.15 | 3.28 | 0.04237 | Decreases | -0.17 |
| **Figure 2B ii.** | | | | | | | | | | |
| **Parameters** | **Conditions** | **Set** | **Dish** | **Well** | **ntotal** | | **Median** | **p-values** | **Trend** | **Fractional change** |
| Base Fluorescence Lifetime (ns) | Control | 3 | 4 | 8 | 57 | | 0.00000000387 | | | |
| | EzrInh | 3 | 4 | 8 | 57 | | 3.62E-09 | 1.56E-16 | Decreasing | -0.06 |
| Mid Fluorescence Lifetime (sec) | Control | 3 | 4 | 8 | 57 | | 4.4E-09 | | | |
| | EzrInh | 3 | 4 | 8 | 57 | | 4.18E-09 | 7.01E-12 | Decreasing | -0.05 |
| **Figure 2B iii.** | | | | | | | | | | |
| **Parameters** | **Conditions** | **Set** | **Dish** | **Well** | **ntotal** | | **Median** | **p-values** | **Trend** | **Fractional change** |
| Fraction of cells Mid-to-base fluorescence lifetime ratio | | | | | | | | | | |
| < 1.125 | Control | 3 | 4 | 8 | 57 | | 1.04E+00 | | | |

| | | | | | | | | | | |
|---|---|---|---|---|---|---|---|---|---|---|
| < 1.125 | EzrInh | 3 | 4 | 8 | 57 | | 1.05E+00 | 6.94E-01 | NS | |
| > 1.125 & < 1.275 | Control | 3 | 4 | 8 | 57 | | 1.18E+00 | | | |
| > 1.125 & < 1.275 | EzrInh | 3 | 4 | 8 | 57 | | 1.20E+00 | 5.65E-01 | NS | |
| >1.275 | Control | 3 | 4 | 8 | 57 | | 1.32E+00 | | | |
| >1.275 | EzrInh | 3 | 4 | 8 | 57 | | 1.40E+00 | 4.57E-02 | Increases | 0.06 |
| **Figure 2B iv.** | | | | | | | | | | |
| **Parameters** | **Conditions** | **Set** | **Dish** | | **ntotal** | | **Median** | **p-values** | **Trend** | **Fractional change** |
| Fluorescence Lifetime (ns) Control | Before HS | 3 | 5 | | 28 | | 4.40E-09 | | | |
| | 5 min | 3 | 5 | | 28 | | 4.57E-09 | 3.88E-04 | Increasing | 0.04 |
| | 20 min | 3 | 5 | | 30 | | 4.42E-09 | 0.26 **wrt Before HS** 0.029 **wrt 5 min** | Decreasing **wrt to 5 min** | -0.03 |
| **Figure 2C ii.** | | | | | | | | | | |
| **Parameters** | **Conditions** | **Set** | | **ntotal** | **Median** | **SEM** | | **p-values** | **Trend** | **Fractional change** |
| Fold change in ratio of edge tension and in tension | | | | | | | | | | |
| Control | Before HS | 4 | | 23 | 1 | 0 | | | | |
| Control | 5 min | 4 | | 23 | 0.91993 | 0.06129 | | | | |

| | | | | | | | | | | |
|---|---|---|---|---|---|---|---|---|---|---|
| Control | 10 min | 4 | | 23 | 1.05 | 0.09155 | | | | |
| Control | 15 min | 4 | | 23 | 1.12 | 0.12898 | | | | |
| Control | 20 min | 4 | | 23 | 1.31448 | 0.29973 | | | | |
| EzrInh | Before HS | 4 | | 24 | 1 | 0 | | | | |
| EzrInh | 5 min | 4 | | 24 | 1.03698 | 0.07002 | | | | |
| EzrInh | 10 min | 4 | | 24 | 1.1066 | 0.08812 | | | | |
| EzrInh | 15 min | 4 | | 24 | 1.00009 | 0.08819 | | | | |
| | 20 min | 4 | | 24 | 1.17016 | 0.08777 | | | | |
| **Figure 2E i.** | | | | | | | | | | |
| **Parameters** | **Conditions** | **Set** | **Dish** | **No. of kymographs** | **ntotal** | **No. of protrusions** | **Median** | **p-values** | **Trend** | **Fractional change** |
| Total no. of Protrusions and retractions per cell | Control | 6 | 13 | 35 | 15 | 52 | 3.00 | | | |
| | EzrInh | 6 | 16 | 63 | 20 | 95 | 3.50 | 0.247 | NS | |
| **Figure 2E iv.** | | | | | | | | | | |
| **Parameters** | **Conditions** | **Set** | **Dish** | **No. of kymographs** | **ntotal** | **No. of protrusions** | **Median** | **p-values** | **Trend** | **Fractional change** |
| **Retraction potential** | | | | | | | | | | |
| Rate of high potential | Control | 6 | 13 | 35 | 15 | | 1.55 | | | |
| | EzrInh | 6 | 16 | 63 | 20 | | 1.08 | 0.02 | Decreases | -0.30 |
| Rate of low potential | Control | 6 | 13 | 35 | 15 | | 0.11 | | | |

| | | | | | | | | | | |
|---|---|---|---|---|---|---|---|---|---|---|
| | EzrInh | 6 | 16 | 63 | 20 | | 0.19 | 0.11 | NS | |
| **Figure 3B ii.** | | | | | | | | | | |
| **Parameters** | **Conditions** | **Set** | **Dish** | | **ntotal** | | **Median** | **p-values** | **Trend** | **Fractional change** |
| Total no. of SF | Control | 3 | 3 | | 169 | | 21 | | | |
| | EzrInh | 3 | 3 | | 175 | | 29 | <0.0001 | Increases | 0.38 |
| Spread area (in $\mu m^2$) | Control | 3 | 3 | | 169 | | 236.3 | | | |
| | EzrInh | 3 | 3 | | 175 | | 446.1 | <0.0001 | Increases | 0.89 |
| **Figure 3D i.** | | | | | | | | | | |
| **Parameters** | **Conditions** | **Set** | **Dish** | | **ntotal** | | **Median** | **p-values** | **Trend** | **Fractional change** |
| Average no. of cortical sf per cell | Control | 3 | 3 | | 169 | | 14 | | | |
| | EzrInh | 3 | 3 | | 175 | | 21 | <0.0001 | Increases | 0.50 |
| **Figure 3D ii.** | | | | | | | | | | |
| **Parameters** | **Conditions** | **Set** | **Dish** | | **ntotal** | | **Median** | **p-values** | **Trend** | **Fractional change** |
| Average no. of peripheral sf per cell | Control | 3 | 3 | | 169 | | 7 | | | |
| | EzrInh | 3 | 3 | | 175 | | 7 | 0.67 | NS | |
| **Figure 3D iii.** | | | | | | | | | | |
| **Parameters** | **Conditions** | **Set** | **Dish** | | **ntotal** | | **Median** | **p-values** | **Trend** | **Fractional change** |
| Mean Intensity of cortical | Control | 3 | 3 | | 169 | | 5.94 | | | |

| | | | | | | | | | | |
|---|---|---|---|---|---|---|---|---|---|---|
| sf per cell | | | | | | | | | | |
| | EzrInh | 3 | 3 | | 175 | | 6.10 | 0.15 | NS | |
| **Figure 3D iv.** | | | | | | | | | | |
| **Parameters** | **Conditions** | **Set** | **Dish** | | **ntotal** | | **Median** | **p-values** | **Trend** | **Fractional change** |
| Mean Intensity of peripheral sf per cell | Control | 3 | 3 | | 169 | | 6.01 | | | |
| | EzrInh | 3 | 3 | | 175 | | 6.52 | 0.05 | NS | |
| **Figure 3E ii.** | | | | | | | | | | |
| **Parameters** | **Conditions** | **Set** | **ntotal** | **Mean (Pa)** | **SD (Pa)** | **Median** | **SEM** | **p-values** | **Trend** | **Fractional change** |
| Maximum traction stress | Before EzrInh | 2 | 152 | 139.00 | 70 | 129 | 6 | | | |
| | After EzrInh | 2 | 152 | 163.00 | 71 | 151 | 6 | 0.00334 | | |
| **Figure 4B i.** | | | | | | | | | | |
| **Parameters** | **Conditions** | **Set** | **Dish** | | **ntotal** | | **Median** | **p-values** | **Trend** | **Fractional change** |
| Spread area (in µm$^2$) | ROCK | 3 | 3 | | 157 | | 421.54 | | | |
| | ROCK+ EzrInh | 3 | 3 | | 172 | | 512.07 | <0.0001 | Increases | 0.21 |
| **Figure 4B ii.** | | | | | | | | | | |
| **Parameters** | **Conditions** | **Set** | **Dish** | | ***n*1** | | **Median** | **p-values** | **Trend** | **Fractional change** |
| Total no. of SF | ROCK | 3 | 3 | | 157 | | 20 | | | |
| | ROCK+ EzrInh | 3 | 3 | | 172 | | 26 | <0.0001 | Increases | 0.30 |
| **Figure 4B iii.** | | | | | | | | | | |

| Parameters | Conditions | Set | Dish | | ntotal | | Median | p-values | Trend | Fractional change |
|---|---|---|---|---|---|---|---|---|---|---|
| Spread area (in μm$^2$) | SMIFH2 | 3 | 3 | | 133 | | 377.52 | | | |
| | SMIFH2 +EzrInh | 3 | 3 | | 94 | | 394.28 | 0.44 | NS | |
| **Figure 4B iv.** | | | | | | | | | | |
| **Parameters** | **Conditions** | **Set** | **Dish** | | **ntotal** | | **Median** | **p-values** | **Trend** | **Fractional change** |
| Total no. of SF | SMIFH2 | 3 | 3 | | 133 | | 24 | | | |
| | SMIFH2 +EzrInh | 3 | 3 | | 94 | | 27 | 0.07 | NS | |
| **Figure 4B v.** | | | | | | | | | | |
| **Parameters** | **Conditions** | **Set** | **Dish** | | **ntotal** | | **Median** | **p-values** | **Trend** | **Fractional change** |
| Average no. of cortical sf per cell | ROCK | 3 | 3 | | 157 | | 13 | | | |
| | ROCK+ EzrInh | 3 | 3 | | 172 | | 16 | <0.0001 | Increases | 0.23 |
| **Figure 4B vi.** | | | | | | | | | | |
| **Parameters** | **Conditions** | **Set** | **Dish** | | **ntotal** | | **Median** | **p-values** | **Trend** | **Fractional change** |
| Average no. of peripheral sf per cell | ROCK | 3 | 3 | | 157 | | 8 | | | |
| | ROCK+ EzrInh | 3 | 3 | | 172 | | 8 | 0.88 | NS | |
| **Figure 4B vii.** | | | | | | | | | | |
| **Parameters** | **Conditions** | **Set** | **Dish** | **Quadrant** | **ntotal** | ***n*2** | **Median** | **p-values** | **Trend** | **Fractional change** |

| | | | | | | | | | | |
|---|---|---|---|---|---|---|---|---|---|---|
| Average no. of cortical sf per cell | SMIFH2 | 3 | 3 | | 133 | | 15 | | | |
| | SMIFH2 +EzrInh | 3 | 3 | | 94 | | 19.50 | <0.0 001 | Increa ses | 0.30 |
| **Figure 4B viii.** | | | | | | | | | | |
| **Parameters** | **Conditions** | **Set** | **Dish** | | **ntotal** | | **Median** | **p-values** | **Trend** | **Fractional change** |
| Average no. of peripheral sf per cell | SMIFH2 | 3 | 3 | | 133 | | 9 | | | |
| | SMIFH2 +EzrInh | 3 | 3 | | 94 | | 6 | <0.0 001 | Decre ases | -0.33 |
| **Figure 4B ix.** | | | | | | | | | | |
| **Parameters** | **Conditions** | **Set** | **Dish** | | **ntotal** | | **Median** | **p-values** | **Trend** | **Fractional change** |
| Mean Intensity of cortical SF (AU) | ROCK | 3 | 3 | | 157 | | 204.93 | <0.0 001 | Decre ases | -0.32 |
| | ROCK+ EzrInh | 3 | 3 | | 172 | | 336.98 | <0.0 001 | Increa ses | 0.64 |
| **Figure 4B x.** | | | | | | | | | | |
| **Parameters** | **Conditions** | **Set** | **Dish** | **Quadrant** | **ntotal** | ***n*2** | **Median** | **p-values** | **Trend** | **Fractional change** |
| Mean Intensity of peripheral SF (AU) | ROCK | 3 | 3 | | 157 | | 207.00 | <0.0 001 | Decre ases | -0.35 |
| | ROCK+ EzrInh | 3 | 3 | | 172 | | 334.00 | <0.0 001 | Increa ses | 0.62 |
| **Figure 4B xi.** | | | | | | | | | | |
| **Parameters** | **Conditions** | **Set** | **Dish** | | **ntotal** | | **Median** | **p-values** | **Trend** | **Fractional change** |

| | | | | | | | | | | |
|---|---|---|---|---|---|---|---|---|---|---|
| Mean Intensity of cortical SF (AU) | SMIFH2 | 3 | 3 | | 133 | | 283.10 | 0.13 | NS | |
| | SMIFH2 +EzrInh | 3 | 3 | | 94 | | 295.49 | 0.02 | NS | |
| **Figure 4B xii.** | | | | | | | | | | |
| **Parameters** | **Conditions** | **Set** | **Dish** | | **ntotal** | | **Median** | **p-values** | **Trend** | **Fractional change** |
| Mean Intensity of peripheral SF (AU) | SMIFH2 | 3 | 3 | | 133 | | 308.94 | 0.33 | NS | |
| | SMIFH2 +EzrInh | 3 | 3 | | 94 | | 342.73 | 0.43 | NS | |
| **Figure 4C ii.** | | | | | | | | | | |
| **Parameters** | **Conditions** | **Set** | **Dish** | **Quadrant** | **ntotal** | **nrup** | **Median** | **p-values** | **Trend** | **Fractional change** |
| Rupture propensity | SMIFH2 | 3 | 5 | 20 | 2910 | 171 | 2.21 | | | |
| | SMIFH2 +EzrInh | 3 | 6 | 24 | 4057 | 8 | 0.00 | <0.0001 | Decreases | -1.00 |
| Rupture Time | SMIFH2 | 3 | 5 | 20 | 2910 | 171 | 388 | | | |
| | SMIFH2 +EzrInh | 3 | 6 | 24 | 4057 | 8 | 53 | 0.002 | Decreases | -0.86 |
| Rupture time categorised (in sec) | | | | | | | | | | |
| <= 300 | SMIFH2 | 3 | 5 | 20 | 2910 | 48 | 226 | 0.03 | | |
| <= 300 | SMIFH2 +EzrInh | 3 | 6 | 24 | 3383 | 6 | 22 | 0.00 | Decreases | -0.90 |
| > 300 & < 600 | SMIFH2 | 3 | 5 | 20 | 2910 | 120 | 426 | 0.66 | NS | |
| > 300 & < 600 | SMIFH2 +EzrInh | 3 | 6 | 24 | 3383 | 2 | 442 | 0.83 | NS | |
| >= 600 | SMIFH2 | 3 | 5 | 20 | 1693 | 3 | 600 | | | |

| | | | | | | | | | | |
|---|---|---|---|---|---|---|---|---|---|---|
| >= 600 | SMIFH2 +EzrInh | 3 | 6 | 24 | NA | NA | | | | |
| % cells ruptured (per quad) | | | | | | | | | | |
| <= 300 | SMIFH2 | 3 | 5 | 13 | 1956 | 48 | 1.60 | | | |
| <= 300 | SMIFH2 +EzrInh | 3 | 6 | 3 | 528 | 6 | 0.65 | 0.23 | NS | |
| > 300 & < 600 | SMIFH2 | 3 | 5 | 15 | 2234 | 120 | 2.70 | | | |
| > 300 & < 600 | SMIFH2 +EzrInh | 3 | 6 | 2 | 327 | 2 | 0.61 | | | |
| >= 600 | SMIFH2 | 3 | 5 | 3 | 423 | 3 | 0.69 | | | |
| >= 600 | SMIFH2 +EzrInh | 3 | 6 | | | | | | | |
| **Figure 4D iii.** | | | | | | | | | | |
| **Parameters** | **Conditions** | **Set** | **Dish** | **No. of kymographs** | **ntotal** | **No. of protrusions** | **Median** | **p-values** | **Trend** | **Fractional change** |
| Total no. of Protrusions and retractions per cell | SMIFH2 | 2 | 7 | 28 | 8 | 8 | 2.50 | | | |
| | SMIFH2 +EzrInh | 2 | 7 | 18 | 7 | 7 | 2.00 | 0.884 | NS | |
| **Figure 4D iv.** | | | | | | | | | | |
| **Parameters** | **Conditions** | **Set** | **Dish** | **No. of kymographs** | **ntotal** | **No. of protrusions** | **Median** | **p-values** | **Trend** | **Fractional change** |
| **Retraction potential** | | | | | | | | | | |
| Rate of high potential | SMIFH2 | 2 | 7 | 28 | 8 | 8 | 1.20 | | | |
| | SMIFH2 +EzrInh | 2 | 7 | 18 | 7 | 7 | 1.59 | 0.28 | NS | |
| Rate of low | SMIFH2 | 2 | 7 | 28 | 8 | 8 | 0.14 | | | |

| | | | | | | | | | |
|---|---|---|---|---|---|---|---|---|---|
| potentia l | | | | | | | | | |
| | SMIFH2 +EzrInh | 2 | 7 | 18 | 7 | 7 | 0.11 | 0.79 | NS |

Table S2

| Figure S1 (B) | | | | | | | | | | |
|---|---|---|---|---|---|---|---|---|---|---|
| **Parameters** | **Conditions** | **Set** | | | **# of events** | | **Median** | **Mean** | **Trend** | **Fractional change** |
| Mean intensity of p-ezr | Secondary control | 1 | | | 754 | | 64.8 | 77.98 | | |
| | Control | 1 | | | 2205 | | 1624.86 | 2360.98 | | |
| | EzrInh | 1 | | | 1112 | | 1773.09 | 2669.21 | | |
| | Secondary control | 2 | | | 744 | | 40.5 | 234.83 | | |
| | Control | 2 | | | 1584 | | 52.65 | 82.17 | | |
| | EzrInh | 2 | | | 2154 | | 63.18 | 131.57 | | |
| | Unstained | 3 | | | 3717 | | 14.11 | 16.96 | | |
| | Secondary control | 3 | | | 6294 | | 19.92 | 27.05 | | |
| | Control | 3 | | | 6177 | | 500.49 | 521.9 | | |
| | EzrInh | 3 | | | 602 | | 19.09 | 262.42 | | |
| **Figure S4 (B)** | | | | | | | | | | |
| **Parameters** | **Conditions** | **Set** | **ntotal** | **Time** | **Mean** | **SD** | **SE** | **Median** | **P-value from 0 min** | |
| Normalized Tension | Control | 4 | 26 | 0 min | 1 | 0 | 0 | 1 | | |
| | Control | 4 | 26 | 2 min | 1.39193 | 0.63753 | 0.13293 | 1.32822 | 0.06029 | |
| | Control | 4 | 26 | 5 min | 1.65613 | 0.87263 | 0.18196 | 1.44952 | 5.364E-05 | |
| | Control | 4 | 26 | 10 min | 2.13512 | 1.04866 | 0.21866 | 1.85287 | 3.064E-07 | |
| | Control | 4 | 26 | 20 min | 2.87736 | 1.69696 | 0.35384 | 2.5842 | 3.064E-07 | |
| | EzrInh | 4 | 29 | 0 min | 1 | 0 | 0 | 1 | | |
| | EzrInh | 4 | 29 | 2 min | 1.27603 | 0.54175 | 0.10625 | 1.20379 | 0.01129 | |
| | EzrInh | 4 | 29 | 5 min | 1.54912 | 0.61829 | 0.12126 | 1.31984 | 2.333E-08 | |

| | | | | | | | | | |
|---|---|---|---|---|---|---|---|---|---|
| | EzrInh | 4 | 29 | 10 min | 1.87682 | 0.70883 | 0.13901 | 1.82565 | 1.102E-09 |
| | EzrInh | 4 | 29 | 20 min | 2.8084 | 3.17847 | 0.62335 | 2.1446 | 2.333E-08 |
| Normalized $SD_{space}$ | Control | 4 | 26 | 0 min | 1 | 0 | 0 | 1 | |
| | Control | 4 | 26 | 2 min | 0.85474 | 0.14686 | 0.03062 | 0.88509 | 5.364E-05 |
| | Control | 4 | 26 | 5 min | 0.85678 | 0.14016 | 0.02923 | 0.86694 | 1.518E-08 |
| | Control | 4 | 26 | 10 min | 0.87797 | 0.13545 | 0.02824 | 0.87057 | 3.064E-07 |
| | Control | 4 | 26 | 20 min | 0.90735 | 0.15793 | 0.03293 | 0.90418 | 0.0004669 |
| | EzrInh | 4 | 29 | 0 min | 1 | 0 | 0 | 1 | |
| | EzrInh | 4 | 29 | 2 min | 0.87315 | 0.18639 | 0.03655 | 0.85539 | 4.92E-05 |
| | EzrInh | 4 | 29 | 5 min | 0.87109 | 0.16539 | 0.03244 | 0.83927 | 0.0003842 |
| | EzrInh | 4 | 29 | 10 min | 0.87467 | 0.22258 | 0.04365 | 0.86776 | 0.00235 |
| | EzrInh | 4 | 29 | 20 min | 0.89136 | 0.18571 | 0.03642 | 0.85473 | 0.01129 |
| Normalized $Sd_{time}$ | Control | 4 | 26 | 0 min | 1 | 0 | 0 | 1 | |
| | Control | 4 | 26 | 2 min | 0.9264 | 0.16304 | 0.034 | 0.89768 | 0.01557 |
| | Control | 4 | 26 | 5 min | 0.85491 | 0.14526 | 0.03029 | 0.8477 | 4.665E-06 |
| | Control | 4 | 26 | 10 min | 0.80847 | 0.12043 | 0.02511 | 0.80939 | 1.518E-08 |
| | Control | 4 | 26 | 20 min | 0.75259 | 0.12417 | 0.02589 | 0.74965 | 5.658E-10 |
| | EzrInh | 4 | 29 | 0 min | 1 | 0 | 0 | 1 | |
| | EzrInh | 4 | 29 | 2 min | 0.97524 | 0.16782 | 0.03291 | 0.9637 | 0.31369 |
| | EzrInh | 4 | 29 | 5 min | 0.89644 | 0.13291 | 0.02607 | 0.92893 | 4.924E-06 |
| | EzrInh | 4 | 29 | 10 min | 0.84462 | 0.13688 | 0.02684 | 0.848 | 4.924E-06 |
| | EzrInh | 4 | 29 | 20 min | 0.73551 | 0.12685 | 0.02488 | 0.72267 | 1.102E-09 |
| Normalized Confinement | Control | 4 | 26 | 0 min | 1 | 0 | 0 | 1 | |

| | | | | | | | | | |
|---|---|---|---|---|---|---|---|---|---|
| | Control | 4 | 26 | 2 min | 1.55014 | 0.77687 | 0.16199 | 1.48069 | 0.01557 |
| | Control | 4 | 26 | 5 min | 2.59481 | 1.98741 | 0.4144 | 1.7399 | 5.364E-05 |
| | Control | 4 | 26 | 10 min | 4.51203 | 5.01506 | 1.04571 | 2.08558 | 0.0004669 |
| | Control | 4 | 26 | 20 min | 9.56068 | 11.4743 | 2.39256 | 4.17254 | 1.518E-08 |
| | EzrInh | 4 | 29 | 0 min | 1 | 0 | 0 | 1 | |
| | EzrInh | 4 | 29 | 2 min | 7304770 | 3E+07 | 5961940 | 1.37964 | 0.01129 |
| | EzrInh | 4 | 29 | 5 min | 1.3E+07 | 5.4E+07 | 1.1E+07 | 2.07313 | 0.0003842 |
| | EzrInh | 4 | 29 | 10 min | 1.1E+08 | 4.9E+08 | 9.5E+07 | 2.61005 | 0.00235 |
| | EzrInh | 4 | 29 | 20 min | 1.2E+08 | 5.1E+08 | 1E+08 | 4.61776 | 2.333E-08 |
| Normalized Active temperature | Control | 4 | 26 | 0 min | 1 | 0 | 0 | 1 | |
| | Control | 4 | 26 | 2 min | 0.98009 | 0.08162 | 0.01702 | 1 | 0.18071 |
| | Control | 4 | 26 | 5 min | 1.01751 | 0.08212 | 0.01712 | 1 | 0.18071 |
| | Control | 4 | 26 | 10 min | 1.02366 | 0.13921 | 0.02903 | 1 | 0.06029 |
| | Control | 4 | 26 | 20 min | 1.09022 | 0.17253 | 0.03597 | 1.00001 | 5.364E-05 |
| | EzrInh | 4 | 29 | 0 min | 1 | 0 | 0 | 1 | |
| | EzrInh | 4 | 29 | 2 min | 1.0438 | 0.28115 | 0.05514 | 1 | 0.03922 |
| | EzrInh | 4 | 29 | 5 min | 0.98101 | 0.11166 | 0.0219 | 1 | 0.0008839 |
| | EzrInh | 4 | 29 | 10 min | 1.04001 | 0.12017 | 0.02357 | 1 | 0.0008842 |
| | EzrInh | 4 | 29 | 20 min | 1.0291 | 0.12287 | 0.0241 | 1 | 3.841E-07 |
| Normalized Viscosity | Control | 4 | 26 | 0 min | 1 | 0 | 0 | 1 | |
| | Control | 4 | 26 | 2 min | 1.58686 | 0.881 | 0.1837 | 1.62925 | 0.00309 |
| | Control | 4 | 26 | 5 min | 1.7483 | 1.187 | 0.24751 | 1.51054 | 0.00309 |
| | Control | 4 | 26 | 10 min | 2.63523 | 1.9826 | 0.4134 | 2.07448 | 4.665E-06 |

| | | | | | | | | | | |
|---|---|---|---|---|---|---|---|---|---|---|
| | Control | 4 | 26 | 20 min | 3.54621 | 2.3645 | 0.49303 | 3.38675 | 4.665E-06 | |
| | EzrInh | 4 | 29 | 0 min | 1 | 0 | 0 | 1 | | |
| | EzrInh | 4 | 29 | 2 min | 1.44771 | 0.73453 | 0.14405 | 1.28705 | 4.92E-05 | |
| | EzrInh | 4 | 29 | 5 min | 1.61651 | 0.83185 | 0.16314 | 1.43386 | 3.841E-07 | |
| | EzrInh | 4 | 29 | 10 min | 2.02425 | 1.08836 | 0.21344 | 1.79935 | 3.841E-07 | |
| | EzrInh | 4 | 29 | 20 min | 2.81288 | 1.8668 | 0.36611 | 2.34498 | 3.841E-07 | |
| **Figure S4 (C)** | | | | | | | | | | |
| **Parameters** | **Conditions** | **Set** | **Dish** | **Well** | **ntotal** | | **Median** | **p-values** | **Trend** | **Fractional change** |
| Fluorescence Lifetime (ns) EzrInh | Before HS | 1 | | | 10 | | 4.30E-09 | | | |
| | 5 min | 1 | | | 10 | | 4.39E-09 | 0.0052 | Increases | 0.02 |
| | 20 min | 1 | | | 10 | | 4.26E-09 | 0.9118 wrt before hs, 0.02881 wrt to 5 min | NS wrt control decreases wrt 5 min | -0.03 |
| **Figure S4 (D)** | | | | | | | | | | |
| **Parameters** | **Conditions** | **Set** | **Dish** | **Time** | **ntotal** | | **Median** | **p-values** | **Trend** | **Fractional change** |
| Ratio of Fluorescence Lifetime at mid and basal plane | Control | 1 | | Before HS | | | 1.05942 | | | |
| | EzrInh | 1 | | Before HS | | | 1.10535 | 0.02881 | Increases | 0.04 |
| | Control | 1 | | 5 min | | | 1.21706 | 0.00389 wrt | Increases | 0.10 |

| | | | | | | | | | | |
|---|---|---|---|---|---|---|---|---|---|---|
| | | | | | | | | before hs | | |
| | EzrInh | 1 | | 5 min | | | 1.25672 | 0.85343 wrt control 5 min, 0.02881 wrt EzrInh before HS | NS wrt control 5 min, increases wrt EzrInh before HS | 0.14 |
| | Control | 1 | | 20 min | | | 1.22817 | | | |
| | EzrInh | 1 | | 20 min | | | 1.15694 | 0.66907 | NS | |
| **Figure S7 (A)** | | | | | | | | | | |
| **Parameters** | **Conditions** | **Set** | **Dish** | | **ntotal** | | **Median** | **p-values** | **Trend** | **Fractional change** |
| Ratio of $SF_{Cort}/SF_{Peri}$ no. | Control | 3 | 3 | | 169 | | 2 | | | |
| | EzrInh | 3 | 3 | | 175 | | 2.88889 | <0.0001 | Increases | 0.44 |
| Ratio of $SF_{Cort}/SF_{Peri}$ mean intensity | Control | 3 | 3 | | 175 | | 0.93713 | | | |
| | EzrInh | 3 | 3 | | 169 | | 0.94999 | 0.40969 | NS | |
| **Figure S7 (B)** | | | | | | | | | | |
| **Parameters** | **Conditions** | **Set** | **ntotal** | **Mean (Pa)** | **SD (Pa)** | **Median** | **SEM** | **p-values** | **Trend** | **Fractional change** |
| Average traction stress(in Pa) | Control | 2 | 152 | 40 | 17 | 37 | 1 | | | |
| | EzrInh | 2 | 152 | 47 | 18 | 45 | 1 | 1.25E-04 | Increases | 0.22 |
| **Figure S9 (A)** | | | | | | | | | | |

| **Parameters** | **Conditions** | **Set** | **Dish** | | **ntotal** | | **Median** | **p-values** | **Trend** | **Fractional change** |
|---|---|---|---|---|---|---|---|---|---|---|
| Spread area (in $\mu m^2$) | Control | 3 | 3 | | 169 | | 236.2893 | | | |
| | ROCK Inh | 3 | 3 | | 157 | | 421.537 | <0.0001 | Increases | 0.78 |
| | SMIFH2 | 3 | 3 | | 133 | | 377.521 | **<0.0001** wrt control **0.0**0142 wrt ROCKInh | **Increases** wrt control **Decreases** wrt ROCKInh | **0.598 -0.104** |
| Total no. of SF | Control | 3 | 3 | | 169 | | 21 | | | |
| | ROCK Inh | 3 | 3 | | 157 | | 20 | 0.30822 | NS | |
| | SMIFH2 | 3 | 3 | | 133 | | 24 | **0.01026** wrt control **0.00134** wrt to ROCKInh | NS wrt control **increases** wrt to ROCKInh | 0.20 |
| **Figure S9 (B)** | | | | | | | | | | |
| **Parameters** | **Conditions** | **Set** | **Dish** | | **ntotal** | | **Median** | **p-values** | **Trend** | **Fractional change** |
| Average no. of cortical sf per cell | Control | 3 | 3 | | 169 | | 14 | | | |
| | ROCK Inh | 3 | 3 | | 157 | | 13 | 0.02725 | NS | |

| | | | | | | | | | | |
|---|---|---|---|---|---|---|---|---|---|---|
| | SMIFH2 | 3 | 3 | | 133 | | 15 | **0.42596** wrt control **0.01262** wrt ROCKInh | NS wrt control NS wrt ROCKInh | |
| Average no. of peripheral sf per cell | Control | 3 | 3 | | 169 | | 7 | | | |
| | ROCK Inh | 3 | 3 | | 157 | | 8 | 0.02811 | NS | |
| | SMIFH2 | 3 | 3 | | 133 | | 9 | **<0.0001** wrt control **0.014** wrt ROCKInh | **Increases** wrt control NS wrt ROCKInh | **0.29** |
| Ratio of $SF_{Cort}$/$SF_{Peri}$ no. | Control | 3 | 3 | | 169 | | 2 | | | |
| | ROCK Inh | 3 | 3 | | 157 | | 1.625 | 1.41E-04 | Decreases | -0.78 |
| | SMIFH2 | 3 | 3 | | 133 | | 1.63636 | **0.0249** wrt control **0.40734** wrt ROCKInh | NS wrt control NS wrt ROCKInh | |
| **Figure S9 (C)** | | | | | | | | | | |
| **Parameters** | **Conditions** | **Set** | **Dish** | | **ntotal** | | **Median** | **p-values** | **Trend** | **Fractional change** |
| Mean Intensity of $SF_{Cort}$ | Control | 3 | 3 | | 169 | | 326.0041 | | | |

| | | | | | | | | | | |
|---|---|---|---|---|---|---|---|---|---|---|
| | ROCK Inh | 3 | 3 | | 157 | | 204.9282 | <0.0001 | Decreases | -0.37 |
| | SMIFH2 | 3 | 3 | | 133 | | 283.1029 | **0.13206** wrt control **<0.0001** wrt ROCKInh | NS wrt control **Increases** wrt ROCKInh | **0.381** |
| Mean Intensity of $SF_{Peri}$ | Control | 3 | 3 | | 169 | | 331.2783 | | | |
| | ROCK Inh | 3 | 3 | | 157 | | 206.5107 | <0.0001 | Decreases | -0.38 |
| | SMIFH2 | 3 | 3 | | 133 | | 308.9368 | **0.33041** wrt control **<0.0001** wrt ROCKInh | NS wrt control **Increases** wrt ROCKInh | **0.496** |
| Ratio of $SF_{Cort}/SF_{Peri}$ mean intensity | Control | 3 | 3 | | 169 | | 0.93713 | | | |
| | ROCK Inh | 3 | 3 | | 157 | | 1.00363 | 1.54E-04 | Increases | 0.07 |
| | SMIFH2 | 3 | 3 | | 133 | | 0.92888 | **0.74839** wrt control **1.23479E-4** wrt ROCKInh | NS wrt control **Decreases** wrt ROCKInh | **-0.074** |
